# Specific exclusion of conjugative plasmids from the gut microflora

**DOI:** 10.64898/2026.08.26.747417

**Authors:** Muhammad Kamruzzaman, Alma Y. Wu, Janani Jeyachandran, Jonathan R. Iredell

## Abstract

The rise of dangerous antimicrobial resistance (AMR), especially to the newer carbapenem antibiotics, is largely driven by the spread of conjugative plasmids between bacteria. These plasmids rapidly disseminate among common gut organisms like *Escherichia coli and Klebsiella pneumoniae*, which together account for around half of lethal sepsis and septic shock. Current AMR control measures, such as surveillance and isolation, are often ineffective, and AMR is often detected for the first time when infection is well established.

Natural plasmid entry-exclusion systems (EES) protect bacterial populations from repeated entry by plasmids that are already established, or indeed by any plasmid with a related cognate EES, and we demonstrate here the exploitation of this mechanism for therapeutic purposes. We combined exclusion genes from three major AMR plasmid types (IncM, IncL, and IncC) into an efficient conjugative plasmid backbone and showed that this engineered ‘probiotic plasmid’ prevents invasive AMR plasmids from entering bacterial populations *in vitro* and *in vivo*, in the mouse gut. This offers a promising new strategy to control invasive AMR plasmids and prevent AMR acquisition in high-risk settings, such as in hospitals or AMR-endemic regions.

**Graphical abstract:** 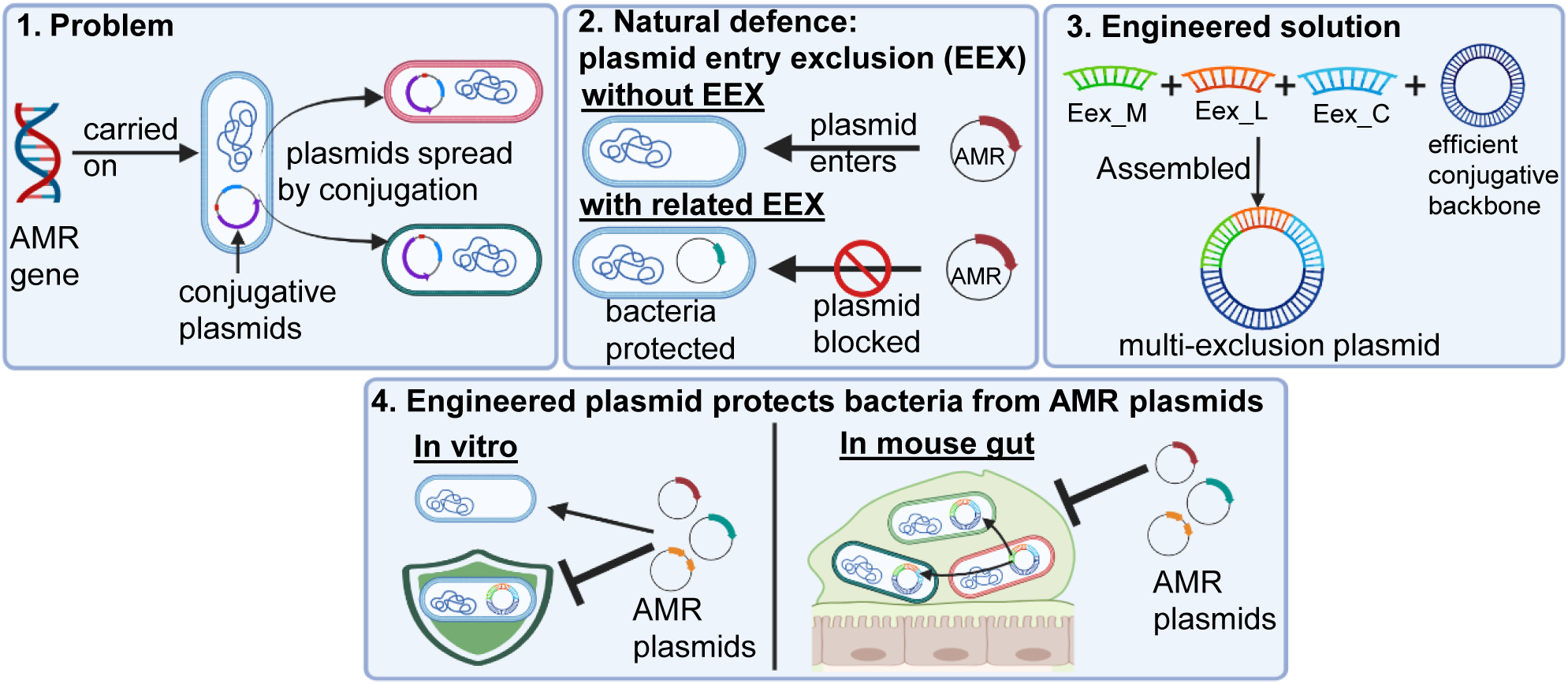

## Introduction

Antimicrobial resistant (AMR) infections are responsible for an estimated five million deaths annually [1–2] and are predicted to be the leading cause of death by 2050 [2–3]. The World Health Organisation (WHO) has recognised the need for urgent, coordinated action to prevent the accelerating spread of antibiotic resistance [4].

A primary driver of this resistance epidemic, particularly in Gram-negative bacteria, is the horizontal transfer of AMR genes across microbial ecosystems, including those of humans, animals, food, and the environment. Primary vehicles are the self-transferable (conjugative) plasmids, highly efficient mobile genetic elements carrying resistance genes for virtually all major antibiotic classes, including extended-spectrum β-lactams, carbapenems, and even ‘last-line’ drugs such as polymixins [5–7]. A significant number of bacterial virulence factors in pathogens like *Salmonella*, *Shigella*, and *E. coli* are also encoded on these plasmids [8].

Bacteria quickly acquire conjugative plasmids from environments in which antibiotic resistance or virulence genes are endemic, including hospitals. Conjugative plasmids efficiently transfer through cell-cell contact [9] and, once acquired, ensure their own persistence in bacterial populations via addiction (typically, toxin-antitoxin) systems that prevent survival of plasmid-free daughter cells [10–11]. In this way, AMR continues to spread and fix in bacterial populations, eroding the value of the antibiotics on which our global health systems and all modern medical advances depend.

Specific entry-exclusion systems (EES) protect recipient bacteria from lethal zygosis due to repeated entry by conjugative plasmids already resident in cells, as these plasmids begin to saturate a population [12–14]. Related plasmids, assigned to Incompatibility (Inc) groups on the basis of interference with each other’s replication [15–16], almost invariably encode entry exclusion proteins (Exc) that interact with cognate targets in the incoming plasmids to directly inhibit transfer after the initial mating pair has formed, typically by 10- to >10,000-fold [12–14, 17–22]. This well-evolved and efficient mechanism therefore provides an attractive opportunity as an anti-AMR strategy.

Antibiotics are expensive to develop [23] and some bacteria are now resistant to all available antibiotics [24]. At present, the strategies for combating AMR spread are limited [25] and new approaches are urgently needed to halt the loss of effective antibiotics. Plasmid curing using different chemical agents [25] and CRISPR-Cas-based approaches [26–28] have been proposed, and direct conjugation inhibitor molecules are being explored for lowering AMR prevalence [29–32], but their *in vivo* efficacy and applicability remained untested. We previously demonstrated efficient target AMR plasmid curing *in vivo* with “conjugative interference plasmids” that efficiently self-propagate within *in vivo* ecosystems, by exploiting replication incompatibility and subverting toxin-antitoxin (‘addiction’) systems to eradicate conjugative AMR plasmids in the gut [33]. This is highly effective in eradicating established plasmids but cannot prevent their first acquisition or later re-acquisition, once cured.

Here, we assembled exclusion genes from IncM, IncL, and IncC plasmids into an efficient conjugative plasmid backbone and showed that it provides a high level of protection to bacteria from acquiring AMR plasmids from IncM, IncL and IncC groups, *in vitro* and *in vivo*.

## Materials and Methods

### Bacterial strains, plasmids, and primers

Bacterial isolates and plasmids (Table 1) and oligonucleotide primers (Supplementary Table S1) used in this study are described. Bacteria were grown in LB-Lennox broth or on LB-Lennox agar at 37°C, and plasmid-carrying bacteria were grown in the presence of the appropriate antibiotic in culture media. J53Az bacteria with the probiotic plasmid were grown and maintained in LB with fosfomycin antibiotic. Bacteria carrying naturally occurring AMR plasmids (IncM, IncL, and IncC plasmids) were grown and maintained in LB with the antibiotic cefotaxime (CTX) and bacteria carrying IncA plasmid (RA1) was grown and maintained in LB with tetracycline (TET). Antibiotics chloramphenicol (CHL, 20 mg/L), sodium azide (AZ, 100 mg/L), rifampicin (RIF, 90 mg/L), fosfomycin (FOF, 200 mg/L), tetracycline (TET, 10 mg/L) and cefotaxime (CTX, 8 mg/L) were used when needed.

**Table 1.** Characteristics of bacterial strains and plasmids used in this study.

| ID | Bacteria /Plasmid | Characteristics | Source/Reference |
| --- | --- | --- | --- |
| DH10B | Bacteria | *Escherichia coli*, used as a host for cloning and recombinant vectors | Cat # EC0113, ThermoFisher Scientific |
| J53Az | Bacteria | Sodium azide resistance *Escherichia coli* K-12 strain | [34] |
| UB5201Rf | Bacteria | Rifampicin resistance derivative of *E. coli* UB5201 strain with *rpoB* mutation S512P | [35] |
| NH78Rf | Bacteria | Rifampicin resistance derivative of *E. coli* ST131 clinical strain with *rpoB* mutation D516G | This study |
| pJIBE401 | Plasmid | Naturally occurring IncM plasmid carrying *bla*<sub>IMP-4</sub> carbapenemase gene | [36] |
| pJIMK45 | Plasmid | pJIBE401 derivative, ~28 kb MRR region replaced with *tetA* gene | [33] |
| pJIMK46 | Plasmid | pJIMK45 derivative, the toxin pemK is replaced with *fosA3* gene | [33] |
| pEc158 | Plasmid | Naturally occurring IncC plasmid carrying *bla*<sub>IMP-4</sub> carbapenemase gene | [37] |
| pJIE1335_L | Plasmid | Naturally occurring IncL plasmid carrying *bla*<sub>OXA-48</sub> carbapenemase gene | [38-39] |
| pExc1_L | Plasmid | *IncL_exc1* gene from pJIE1335_L with its own promoter and RBS was cloned into the XbaI and HindIII sites of the cloning vector pACYC184. | This Study |
| pExc1_M | Plasmid | *IncM_exc1* gene from pJIBE401 with its own promoter and RBS was cloned into XbaI and HindIII sites of the cloning vector pACYC184. | This Study |
| pExc4_L | Plasmid | The synthesised *IncL_exc4* gene with its own promoter and RBS was cloned into the XbaI and HindIII sites of the cloning vector pACYC184. | This Study |
| pExc4_M | Plasmid | The synthesised *IncM_exc4* gene with its own promoter and RBS was cloned into the XbaI and HindIII sites of the cloning vector pACYC184 | This Study |
| pPB1 | Plasmid | Probiotic plasmid, resistance to fosfomycin. The *tetA* gene of pJIMK45 was replaced with *IncL_exc-IncC_exc-fosA3* gBlock | This study |
| pPB1.1 | Plasmid | Probiotic plasmid, resistance to fosfomycin and tetracycline. The *pemK* toxin gene of PB1 was replaced with *tetA* gene | This study |
| pACYC184 | Plasmid | Low copy number (10-12 copies/cell) cloning vector, chloramphenicol and tetracycline resistance | New England Biolabs, USA |
| pKM200 | Plasmid | Lambda red recombinase plasmid, chloramphenicol resistance | [40] |

### Sequence analysis of Exc variants

To identify the plasmid exclusion gene (*exc*) variants present in the plasmid sequences in the GenBank, we performed a BLASTn search with each of the *exc_C*, *exc_L*, and *exc_M* nucleotide sequences from the plasmids pEc158 (Accession no. KY887596), pOXA-48 (Accession no. CP066515) and pEl1573 (Accession no. JX101693), respectively, against the GenBank standard nucleotide sequence database using default parameters, with a maximum target sequence number set to 5000. We downloaded the FASTA file of the aligned sequences and removed any sequences with less than 100% length coverage. Any sequence oriented in reverse order has been made reverse complement using Geneious software (Geneious Prime® 2025.1.3). The entire sequence file was then translated using Geneious, and non-identical amino acid sequences were identified using Geneious or BioEdit (BioEdit 7.2).

### Construction of mini exclusion plasmids

Mini exclusion plasmids for IncM and IncL groups were generated by cloning the corresponding plasmid exclusion gene into the cloning vector pACYC184. The *exc_M* and *exc_L* genes, including their native promoter and ribosome binding site, were PCR-amplified from the IncM plasmid pJIBE401 and the IncL plasmid pJIE1335_L, respectively, using primers listed in Table S1. The forward and reverse primers were designed to incorporate XbaI and HindIII restriction sites, respectively.

PCR amplification was carried out using Phusion High-Fidelity DNA Polymerase (New England Biolabs, USA) in accordance with the manufacturer’s protocol. The resulting PCR products were digested with XbaI and HindIII and independently ligated into similarly restricted pACYC184, a medium-copy-number plasmid, to generate the mini exclusion constructs pExc_M and pExc_L (Table 1). In addition, variant exclusion genes, *IncM_Exc4* and *IncL_Exc4*, were synthesised commercially (Integrated DNA Technologies, USA) with their native promoter and ribosome binding site, as well as flanking XbaI and HindIII restriction sites. These synthetic fragments were cloned into pACYC184 to construct the mini exclusion plasmids pExc4_M and pExc4_L. All constructs were subsequently verified by DNA sequencing to confirm the accuracy of the cloned sequences.

### Liquid mating conjugation

Liquid mating was used to measure plasmid conjugation frequency. *E. coli* NH78Rf with IncM plasmid pJIBE401, IncL plasmid pJIE1335_L, IncA plasmid RA1 or IncC plasmid pEc158 were used as donor bacteria, and sodium azide-resistant bacteria J53Az and J53Az carrying mini exclusion plasmids or the conjugative probiotic exclusion plasmid were used as the recipient for each plasmid transfer. Briefly, donor and recipient bacteria were grown overnight in 10 mL of LB-Lennox broth with the appropriate antibiotics. Cultures were washed twice with an equivalent volume of sterile saline solution (0.85% NaCl) and resuspended in 10 mL LB_Lennox broth. The OD_600_ of each culture was adjusted to 1.5, and 1 mL of each donor and recipient culture was mixed, centrifuged, and the cell pellet was dissolved in 100 µL of saline and transferred into 5 mL LB in a 15 mL Falcon tube. Mating mixtures were incubated at 37°C for 20 h without shaking. Mating was ended by vortexing the mixtures for 10 sec. Mating mixtures were then serially diluted with saline and plated onto LB-Lennox agar plates containing AZ (100 mg/mL) and CTX (8 mg/mL) to select transconjugants for IncM, IncL and IncC plasmids and plated onto agar plates containing AZ (100 mg/mL) and TET (10 mg/mL) plates for IncA transconjugants. Conjugation frequency was calculated by dividing the total number of transconjugants by the number of recipient bacteria added to the conjugation mixture.

### Construction of conjugative probiotic plasmid

The homologous recombination-based allelic exchange method [40] was used to insert the gBlock of *excL-excC-fosA3* fusion gene fragment into the pJIMK45 [33] by replacing the tetracycline resistance marker gene. The *E. coli* UB5201Rf strain carrying pJIMK45 was transformed with a lambda red recombinase plasmid pKM200 (CHL^R^) and selected on TET plus CHL plates at 30°C. Electro-competent cells prepared from UB5201Rf (pJIMK45 + pKM200) were transformed with the PCR-amplified gBlock (∼1.5 µg) of *excL-excC-fosA3* by electroporation and transformants selected on FOF-resistant plates. Selected transformants were then confirmed for sensitivity to TET, and the correct insertion was confirmed by PCR and sequencing.

### Bacterial growth kinetics assay

Growth kinetics of *E. coli* bacteria J53Az and J53Az with different plasmids (probiotic and naturally occurring AMR plasmids) were measured using a SpectrMax plate reader, as described previously [41]. Briefly, a single colony of each strain was grown overnight in LB Lennox broth with the appropriate antibiotics, then subcultured at 1:1000 into 10 mL of LB Lennox broth and grown for 3 hrs. Cultures were then adjusted to 0.5 McFarland using a Nephelometer and diluted 100-fold in LB Lennox medium with appropriate antibiotics. 150 µL of diluted bacterial cultures were transferred in triplicate into a 96-well microplate (Corning, USA) and incubated with shaking at 37°C overnight in a SpectraMax iD5 multimode microplate reader (Molecular Devices, USA). The OD_600_ was measured every 10 min for 16 h. A mean of nine data points, three technical replicates from each of three biological replicates, was used to generate growth curves with standard errors.

### Expression analysis of *exc* genes

The relative expression of *exc* genes (*exc_M*, *exc_L* and *exc_C*) from the constructed probiotic plasmid pPB1.1 was measured using quantitative real-time PCR (qRT-PCR) as described previously [14]. Briefly, *E. coli* J53Az strains with or without probiotic plasmids were grown O/N with appropriate antibiotics. One mL of the stationary phase cultures was harvested by centrifugation at 10,000 rpm for 10 min. Bacterial pellets were treated for 5 min with 400 mg/mL Chicken egg lysozyme (Sigma Aldrich, USA) in TE (10 mM Tris-Cl and 1 mM EDTA) buffer. Total RNA was purified using the NucleoSpin RNA Plus kit (Macherey-Nagel, Germany) according to the manufacturer’s instructions. The total RNA of each sample was treated for 2 h at 37°C with 1 µL of Turbo DNase (Invitrogen, USA) to eliminate genomic DNA. One tenth volume of DNase inactivation reagent (Invitrogen, USA) was added and incubated 5 min at room temperature to inactivate DNase and clear RNA solutions were separated after centrifugation at 11,000 x g for 2 min. Gene-specific PCR was used to confirm the absence of DNA contamination.

Reverse transcription was carried out using one µg of each RNA sample as a template using the High-Capacity cDNA Reverse transcription kit (Applied Biosystems, USA) according to the manufacturer’s instructions. All cDNA samples were amplified using gene-specific qRT-PCR primers (**Table S1**) and SYBR Green PCR reagent (Qiagen, Germany) on a Rotor-Gene 6000 real-time thermocycler. The qPCR program includes initial denaturation at 95°C for 5 min, then 40 cycles of denaturation 95°C for 5 sec and annealing/extension at 60°C for 10 sec.

The CT values of all reactions were obtained using Rotor-Gene Q Series Software. The internal reference gene *rpoB* was used. Using the 2^-ΔΔCT^ method, the relative expression levels of the target gene were normalised to the internal reference gene. The procedure was repeated with four biological replicates of total RNA.

### Mouse model experiment for AMR protection

Five-week-old female BALB/c mice (Animal Resource Centre, Perth, WA, Australia) were used to test the probiotic plasmid for mouse gut microbiota protection from AMR plasmid acquisition. All research and animal care procedures were approved by the Animal Ethics Committee of the Western Sydney Local Health District (animal ethics protocol 4382.02.23) in accordance with the “Australian Code of Practice for the Care and Use of Animals for Scientific Purposes”. Experiments were performed in the Biohazard room of the Westmead Bioresources Facility, and mice were housed in open-lid M1 polypropylene cages (Able Scientific, Australia) on a 12 h light/dark cycle, with food and water available *ad libitum*.

## Results

### Analysis of entry exclusion protein sequences in IncM, L, A and C plasmid types

Plasmid entry exclusion genes have been identified or predicted for most major Inc types of plasmids, but the extent of conservation within an Inc type remains poorly defined. Here, we used BLASTn searches against GenBank sequences with the IncC, IncM, and IncL exclusion genes *exc_C*, *exc_M*, and *exc_L*, respectively, and converted the matched sequences into amino acids to identify Exc variants within each plasmid group.

We identified a single variant of the Exc protein in all IncC plasmids in the GenBank (as of 23^rd^ November 2025). IncC_Exc nucleotide sequence search identified 1296 plasmids in the GenBank, among them 1198 were IncC plasmids, and all had a single Exc variant. The rest of the 98 plasmids were from the IncA group and are a variant of IncC_Exc (Supplementary **Fig. S1A**). IncC_Exc has ∼86% amino acid sequence identity with the IncA_Exc proteins, but it was demonstrated that the Exc_C in recipient bacteria can inhibit acquisition of IncA plasmids by conjugation [13].

BLAST search identified four variants of IncM Exc proteins in the plasmid sequences available in GenBank (**Fig. S1B**). IncM_Exc1 (110/314 hits), IncM_Exc2 (27/314 hits, 52V>I) and IncM_Exc3 (148/314 hits, 52V>I,72A>S), together account for >91% of all IncM plasmids, and there are only one or two neutral amino acid changes among these variants. The rest of the 9% IncM plasmids have the IncM_Exc4 variant (29/314 hits, 35L>I, 171I>L, 207T>S) and also carry only three neutral amino acid differences from the Exc_1 variant.

Within 5 IncL_Exc variants, more than 91% of the IncL plasmids (531/588) in the GenBank have one major variant of Exc (IncL_Exc1 in **Fig. S1C**), and 7% (39/588) have three minor variants of Exc (IncL_Exc2, IncL_Exc3, IncL_Exc5). IncL_Exc2 (I159L), IncL_Exc3 (I159L, A128V) and IncL_Exc_5 (A55T, I159L) have only one and two neutral amino acid changes from the dominant Exc1 type and 1.7% of IncL plasmids have IncL_Exc4 variant (46G>S, 61A>V, 64V>I, 159I>L, 188H>N, 205T>N) with multiple amino acid differences but all are minor and neutral amino acid variations.

IncL and IncM plasmids are related [42–43] but encode distinct Exc proteins with only 35% amino acid sequence identity (IncM_Exc1 Vs IncL_Exc1, **Fig. S1D**).

We predicted the promoter sequences associated with the major Exc variants of IncL and IncM plasmids. All IncL_Exc variants have an identical promoter sequence (Supplementary **Fig. S2A**), and IncM_Exc variants have identical −35 and −10 regions, but 1-2 nucleotide differences were identified in the promoter spacer regions (Supplementary **Fig. S2B**). There is only 35% aa sequence identity between IncL and IncM Exc proteins but their encoding genes possess near-identical promoter sequences (Supplementary **Fig. S2B**).

### Evaluating plasmid exclusion properties of IncM and IncL plasmid-encoded entry-exclusion proteins

The *exc* genes of IncM and IncL plasmids have been predicted based on sequence identity, but their role in inhibiting IncM and IncL plasmid conjugation transfer has not been experimentally demonstrated. Here we cloned major *exc* variants with their natural promoter and ribosome binding site (RBS) from IncM (*IncM_exc1*) and IncL (*IncL_exc1*) plasmids separately into a low copy cloning vector, pACYC184, to construct mini exclusion plasmid pExc1_M and pExc1_L (**Table 1**). We then transformed mini exclusion plasmids into *E. coli* bacteria J53Az to construct J53Az (pExc1_M) and J53Az (pExc1_L) strains. Conjugation transfer inhibition by these cloned *exc* gene products were studied by liquid conjugation mating experiments. Clinical ST131 *E. coli* donor bacteria NH78Rf carrying pJIBE401 (IncM) or pJIE1335_L (IncL) AMR plasmid were mated in liquid LB media with the recipient J53Az carrying either empty vector pACYC184 or one of the mini exclusion plasmids (pExc1_M or pExc1_L). Transconjugants were selected on selective agar plates and conjugation frequency calculated by dividing the number of transconjugants by the number of recipient bacteria. In this experiment, the variable factor is the recipient bacteria, which carry either an empty vector or a mini exclusion plasmid; therefore, instead of the donor, we used the recipient numbers to calculate the transconjugant numbers. Conjugation transfer of both IncM and IncL AMR plasmids was strongly inhibited in the recipient bacteria carrying mini exclusion plasmids with the respective *exc* genes (**Fig. 1A**). This strongly suggests that *IncM_exc* and *IncL_exc* genes can be used to prevent bacteria from acquiring respective AMR plasmids.

**Fig. 1.**
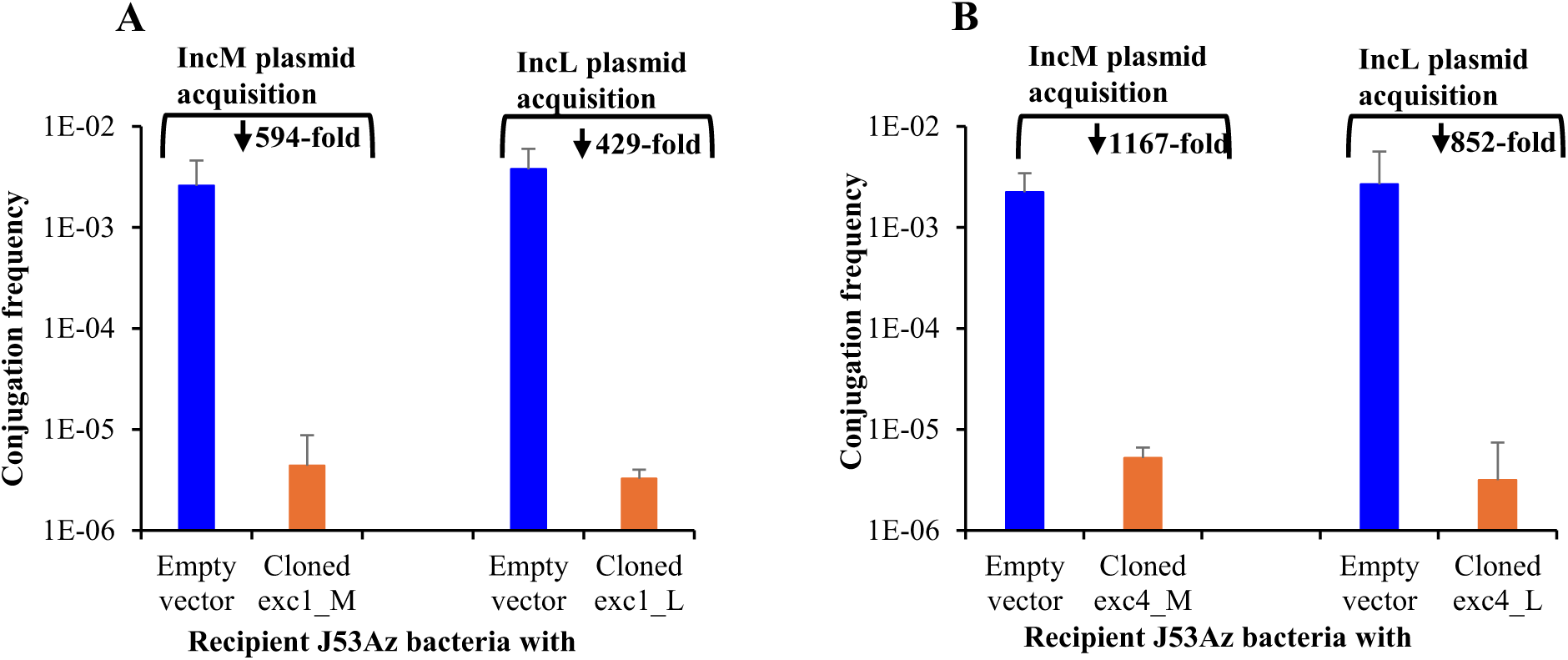
IncM and IncL plasmid conjugation transfer inhibition from the donor NH78Rf to the recipient bacteria J53Az carrying either an empty vector or cloned *exc1_M* or *exc1_L* (**A**) or *exc4_M* or *exc4_L* (**B**). Data is the mean of four independent 6 h liquid mating experiments with standard errors.

### Exc variants do not influence the ability to inhibit conjugation of IncM and IncL plasmids

We therefore investigated whether a single Exc variant is sufficient to protect bacteria against acquisition of all IncM and IncL plasmids. Sequence comparisons showed that Exc variants from IncM and IncL plasmids differ only by a small number of minor or neutral amino acid substitutions, which are unlikely to affect conjugation inhibition across variants.

To test this, we synthesised the IncM_Exc4 and IncL_Exc4 gene variants (Supplementary **Fig. S3**) and cloned them into pACYC184 to generate the mini-exclusion plasmids pExc4_M and pExc4_L, respectively. These variants exhibit the greatest divergence from the Exc1 variants **(Fig. S1B, C**), differing by three amino acids in IncM_Exc4 and six amino acids in IncL_Exc4. The constructs were introduced into J53Az to create J53Az (pExc4_M) and J53Az (pExc4_L) strains.

Liquid mating assays were then performed using donor strain NH78Rf carrying either pJIBE401 (carrying IncM_Exc1 variant) or pJIE1335_L (carrying IncL_Exc1 variant), with recipients containing either the empty vector pACYC184 or the corresponding mini-exclusion plasmid with Exc4 variant. The results showed that recipient bacteria carrying mini-exclusion plasmids with Exc4 variants inhibited the acquisition of IncM and IncL plasmids with Exc1 variants even higher extent than the recipient carrying mini-exclusion plasmid with Exc1 variants (**Fig. 1B**).

These findings strongly indicate that the presence of any Exc variant in a recipient cell is sufficient to exclude the acquisition of all other IncM or IncL plasmid variants.

### Construction of the conjugative probiotic plasmid pPB1.1 to prevent invasion by IncM, IncL, IncC, and IncA plasmids

We previously demonstrated that the IncM plasmid pJIBE401 exhibits highly efficient *in vivo* conjugative transfer, which we exploited to develop AMR curing/interference plasmids for eliminating target plasmids [33]. Building on this system, we used the pJIBE401 backbone pJIMK45, to construct a probiotic plasmid designed to protect bacteria against invasion by four clinically relevant AMR plasmid types. pJIMK45 is the derivative of pJIBE401, where the multi-resistance region and associated transposable elements in pJIBE401 were deleted and replaced with a tetracycline resistance gene (*tetA*) [33].

The starting plasmid pJIMK45 inherently carries the *exc* gene (Exc1 variant) specific to the IncM plasmid type. Therefore, the full-length *exc* genes from IncL and IncC plasmids, including their native promoter and ribosome-binding site, were combined with the *fosA3* gene (conferring resistance to fosfomycin, FOF) (Supplementary **Fig. S4**). This gBlock was synthesised commercially (IDT, USA). Although the resulting gBlock contained a 44 bp deletion within the intergenic region between *excL* and *excC* (Supplementary **Fig. S4A**), the genes and the promoter regions remained intact. Therefore, the gBlock was used as a PCR template to amplify the *excL-excC-fosA3* fragment using Phusion High-Fidelity DNA polymerase with primers EXC1-F and FosA3-R, each containing 40 bp homology extensions matching the insertion site in pJIMK45.

The amplified fragment was used to replace the *tetA* gene in pJIMK45 via homologous recombination, generating the conjugative probiotic plasmid pPB1 (**Fig. 2**), which became resistance to fosfomycin but susceptible to tetracycline. Successful recombinants were verified by tetracycline sensitivity, PCR screening, and Sanger sequencing.

**Fig. 2.**
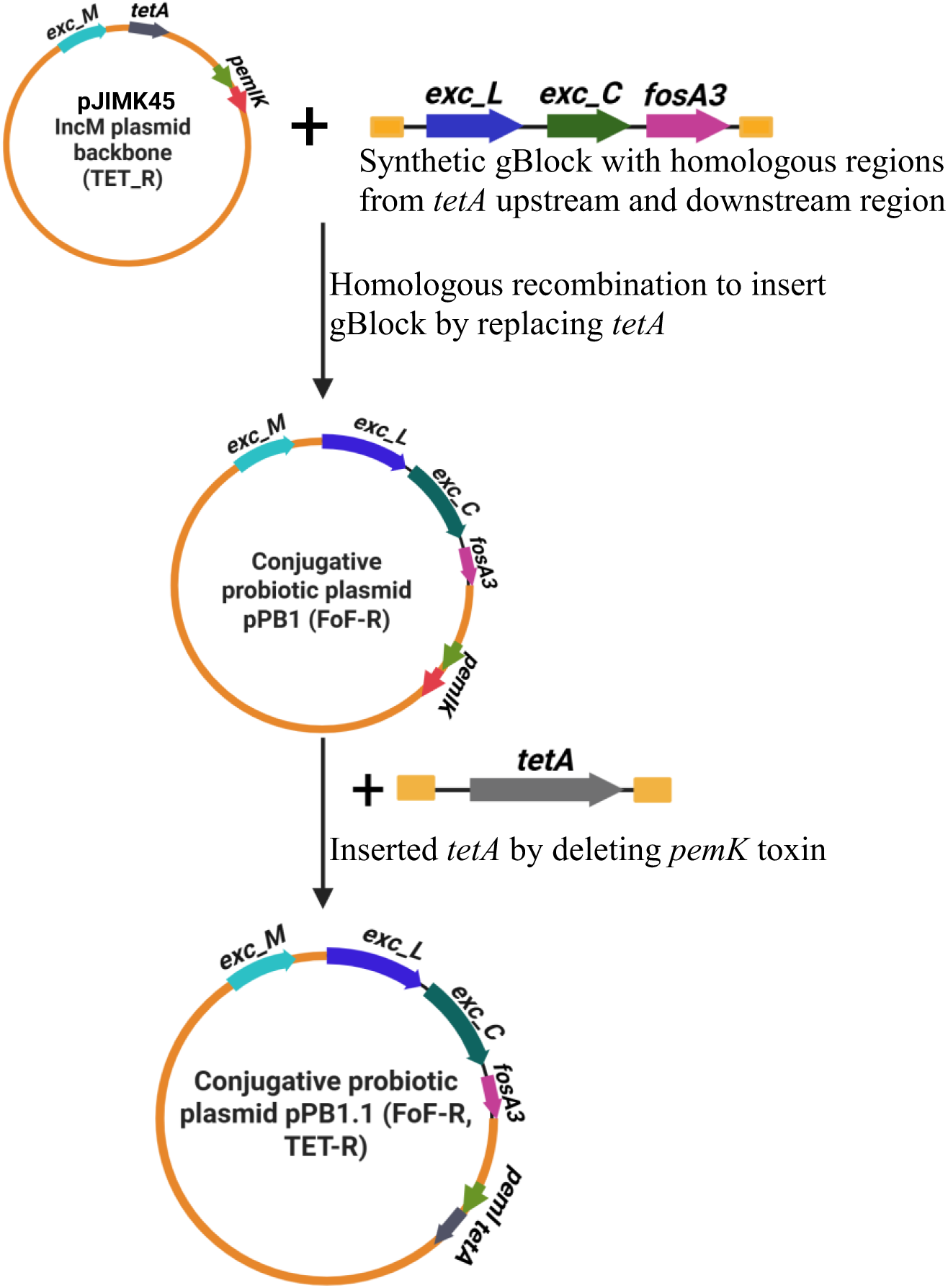
Construction of conjugative probiotic plasmid pPB1.1. Starting plasmid pJIMK45 is the ∼28.5-kb multi-resistance region (MRR) deletion derivative of the natural IncM plasmid pJIBE401. The homologous region of the flanking gBlock and *tetA* genes are shown as yellow bars. This figure was made using BioRender.

The resulting conjugative probiotic plasmid pPB1 retained an intact *pemIK* toxin–antitoxin system, ensuring stable maintenance within bacterial populations. To generate an unstable version of the plasmid, the *pemK* toxin gene was replaced with *tetA*, resulting in plasmid pPB1.1 (**Fig. 2**). A schematic map of pPB1.1 is provided in supplementary **Fig. S5**.

### Sequence analysis of the constructed probiotic plasmid pPB1.1

Sequence analysis of the engineered probiotic plasmid pPB1.1 was performed using Illumina short-read sequencing of UB5201Rf (pPB1.1), which identified three mutations within the construct (Supplementary **Fig. S6**). One single nucleotide polymorphism (A>G) in *tetA* was synonymous and did not alter the amino acid sequence or tetracycline resistance phenotype. A second SNP (G>A) located at the terminal region of *exc_L* resulted in a minor amino acid substitution (G>S), which is predicted to have little or no impact on Exc_L function. Additionally, a single nucleotide deletion (A) was detected in the promoter spacer region of *exc_C*, which is predicted to influence the expression of *exc_C* from the plasmid and has been tested.

### Expression of *exc* genes from the pPB1.1 in *E. coli* bacteria

Specific quantitative real-time PCR assay [41, 44] showed expression of *exc_M* to be very similar to that from the native IncM plasmid, and *exc_L* and *exc_C* to be more strongly expressed from pPB1.1 than from their native plasmids (**Fig. 3, A-C**). The promoter spacer for the *exc_C* in the native plasmid has 22 nucleotides compared to 21 bases in pPB1.1, perhaps contributing to higher-level expression of *exc_C*.

**Fig 3.**
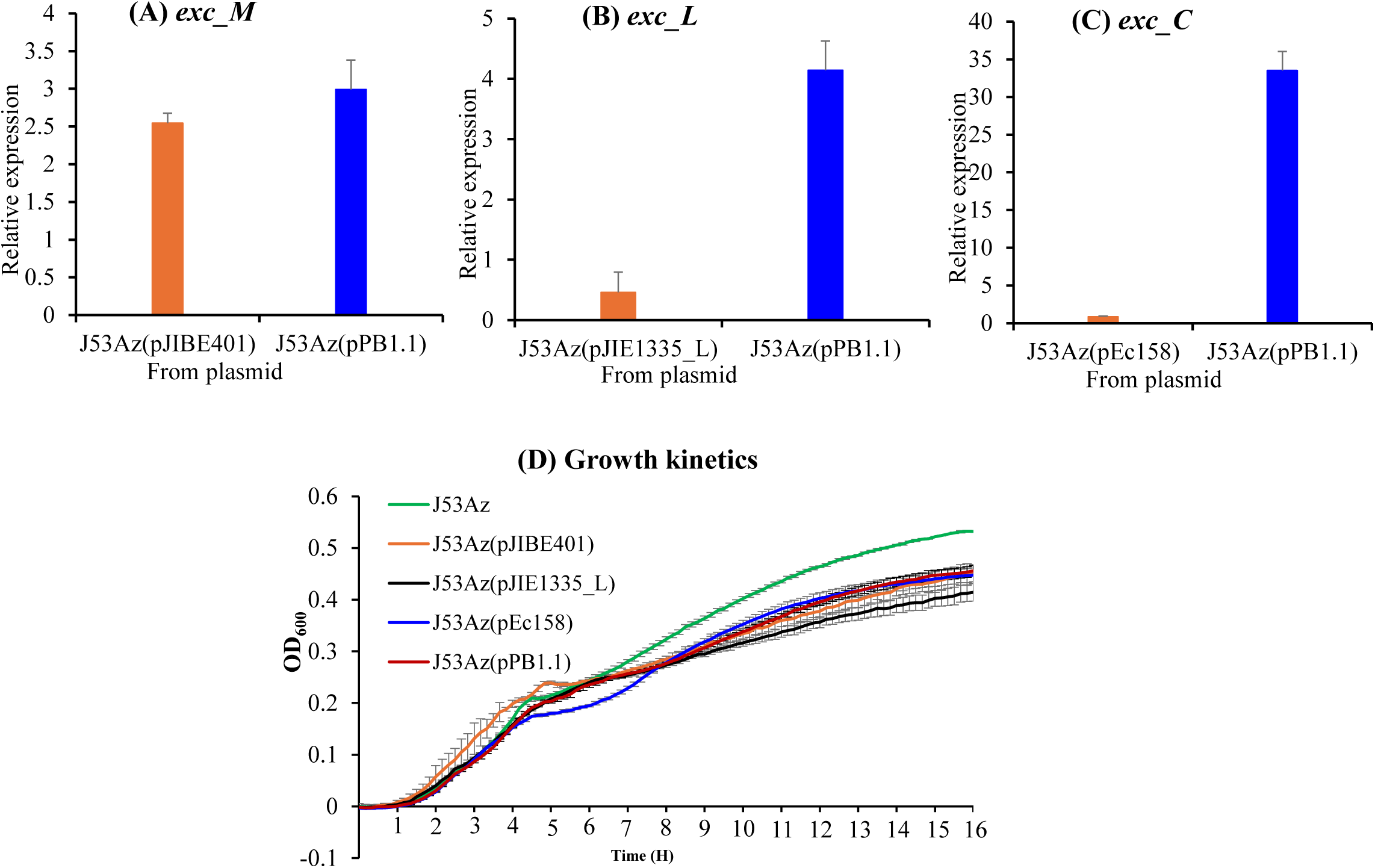
Relative expression of exclusion (*exc*) genes (A-C) and the bacterial fitness cost in growth (D) from the conjugative probiotic plasmid and naturally occurring AMR plasmids. Orange bars are the expression from native AMR plasmids and blue bars are for the expression from the probiotic plasmid pPB1.1. Data are the mean of four independent experiments with standard errors.

### Fitness cost of the probiotic plasmid pPB1.1 in *E. coli*

We investigated whether high-level constitutive expression of *exc_L* and *exc_C* in pPB1.1 imposes an additional fitness burden. Growth kinetics assays were conducted using the host strain *E. coli* J53Az, in the presence or absence of the probiotic plasmid and the naturally occurring AMR plasmid pJIBE401 (the backbone for pPB1.1). We also tested J53Az carrying natural IncL (pJIE1335_L) and IncC (pEc158) plasmids, which are the original sources of *exc_L* and *exc_C*, respectively.

While plasmid-free J53Az showed slightly improved growth, strains harbouring pPB1.1 or the natural IncM, IncL, or IncC plasmids exhibited comparable growth kinetics (**Fig. 3D**). These findings indicate that the engineered probiotic plasmid, despite containing multiple exclusion genes expressed at high levels, does not confer an additional in vitro fitness cost.

### Probiotic plasmid pPB1.1 prevents invasion by IncM, IncL, IncC, and IncA plasmids *in vitro*

We investigated whether acquisition of antimicrobial resistance (AMR) plasmids could be inhibited in bacteria carrying the probiotic plasmid pPB1.1. Liquid mating experiments were conducted between the donor strain, clinical *E. coli* ST131 NH78Rf harboring AMR plasmids from the IncM, IncL, IncC, and IncA groups, and the recipient strain *E. coli* J53Az, either with or without pPB1.1.

In 20-hour mating assays, IncM, IncL, IncC, and IncA plasmids readily transferred into J53Az. In contrast, their transfer into J53Az carrying pPB1.1 was reduced by more than 99.9% (**Fig. 4**). These results demonstrate that pPB1.1 effectively blocks the acquisition of IncM, IncL, IncC, and IncA plasmids.

**Fig 4.**
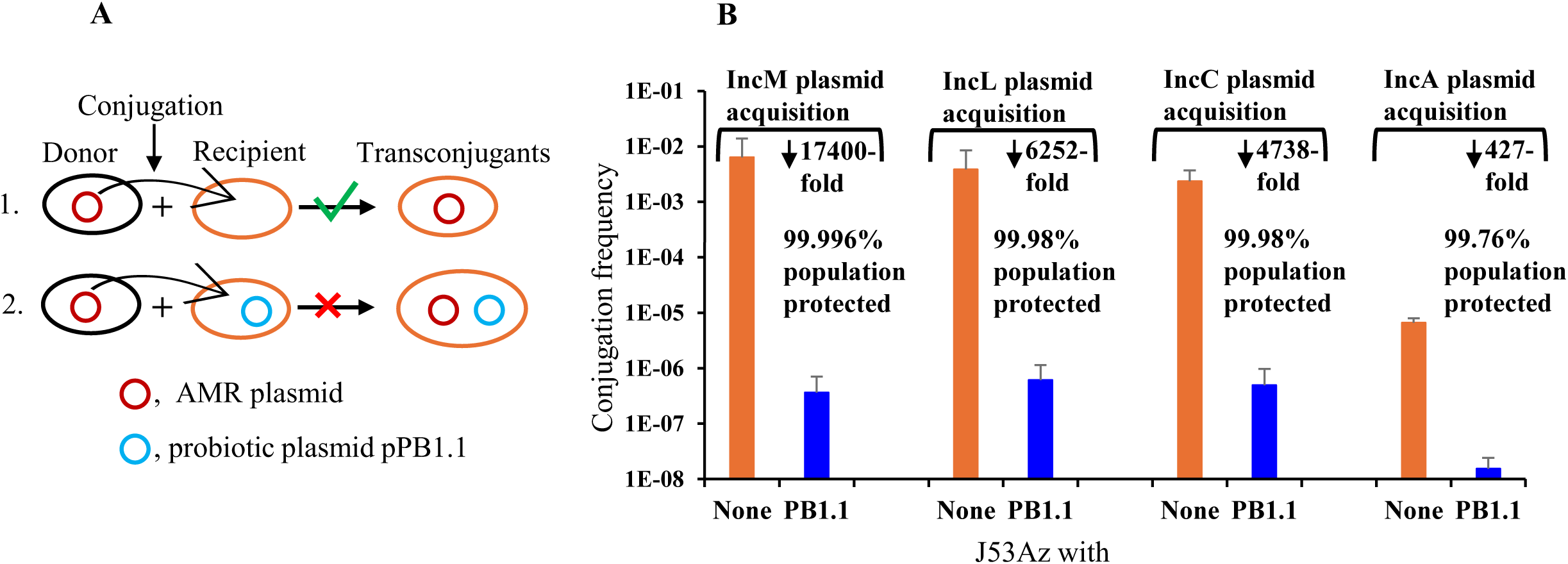
Conjugative probiotic plasmid (pPB1.1) protects bacteria from the acquisition of problem AMR plasmids *in vitro*. (**A**) Liquid mating conjugation assay, from which we measured the number of transconjugants per recipient. Red and blue circles denote the AMR and the probiotic plasmid, respectively. Probiotic plasmid in the recipient bacteria protects them from the acquisition of AMR plasmids. (**B**) Orange bars are for the conjugation frequency of different AMR plasmids into the empty J53Az bacteria during liquid mating experiments. Blue bars are for the conjugation frequency of the respective plasmids into J53Az carrying conjugative probiotic plasmid pPB1.1. The transfer of AMR plasmids is significantly reduced when recipient bacteria carry a probiotic plasmid. Conjugation frequency is the ratio of transconjugants per recipient. Data are the mean of 4 independent liquid mating experiments with standard errors.

### Evaluation of probiotic plasmid pPB1.1 for protection of mouse gut microbiota against AMR plasmid acquisition

We assessed the ability of the conjugative probiotic plasmid pPB1.1 to protect mouse gut microbiota from acquiring AMR plasmids of the IncM, IncL, and IncC types (**Table 1**) during incidental exposure to infected faeces via coprophagy. The experimental protocol including mouse grouping, strains and plasmids used, and sample collection and analysis information are summarised in **Figure 5**.

**Fig. 5.**
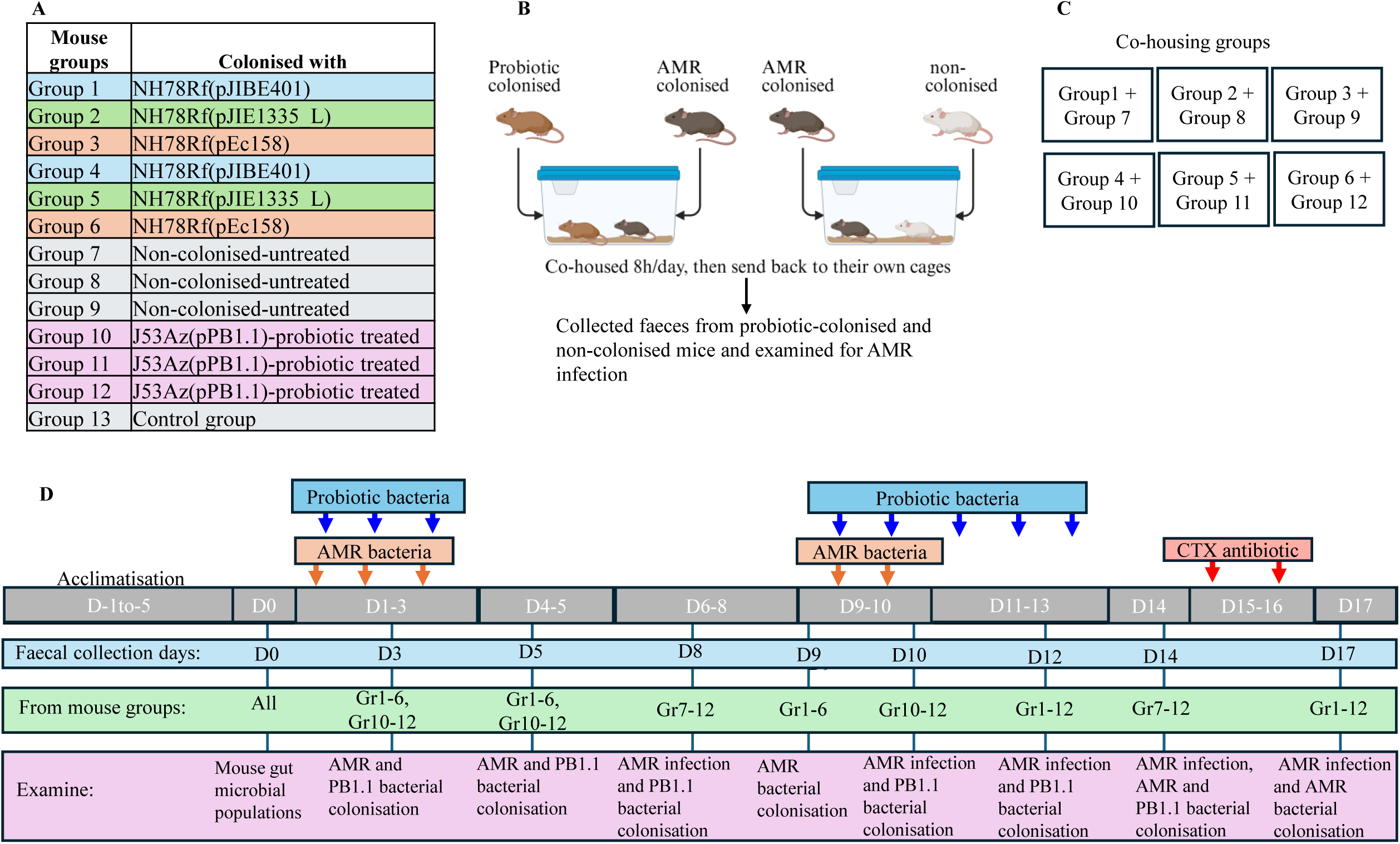
Mouse grouping and colonisation details (A), mouse experimental protocol summary (B), co-housing groups (C), and details of the mouse experimental protocol (D). Mouse gut was colonised with *E. coli* carrying an AMR plasmid or the probiotic plasmid by providing the bacteria in drinking sucrose water. A total of 13 groups of mice were used to test the protection of the mouse gut microbiota against the acquisition of 3 different AMR plasmids (IncM, IncL, and IncC types) using a single probiotic plasmid, pPB1.1 (**A, D**). (**C**) Each box represents which mouse groups are co-housed. (**D**) Orange vertical arrows are for the doses of AMR bacteria, blue arrows are for probiotic bacteria, and red arrows are for CTX antibiotic doses. All of these are provided in a drinking water bottle with 8% sucrose water for O/N. The fresh faecal collection dates, samples collected from which mouse groups and the testing details are mentioned (**D**).

To evaluate this, a total of 13 groups of mice (3 mice per group) were used. One group (Gr13) served as environmental and food control. Six groups (Gr1–6) were colonised with *E. coli* NH78Rf carrying AMR plasmids, with two groups assigned per plasmid type (IncM: Gr1 and Gr4; IncL: Gr2 and Gr5; IncC: Gr3 and Gr6). Three probiotic-treated groups (Gr10–12) were colonised with *E. coli* J53Az carrying the probiotic plasmid pPB1.1, while three untreated groups (Gr7–9) were left uncolonised and all received food and water *ad libitum*.

Upon arrival, mice were acclimatised for five days (days −5 to −1) in standard filter-top cages with free access to food and water. On day 0, prior to bacterial administration, mice were marked with permanent markers, and faecal samples were collected to characterise the native gut microbiota.

Groups 1–6 received *E. coli* carrying AMR plasmids via sucrose water overnight on days 1–3 and 9–10, while groups 10–12 received *E. coli* carrying pPB1.1 overnight on days 1–3 and 9–13. Co-housing experiments were established from day-3: one AMR-colonised group was paired with a probiotic-treated group, and another AMR-colonised group was paired with an untreated group (details in **Fig. 5A-C**). Mice were co-housed for 8 hours per day and separated for the remainder of the time, with access to either standard food and water or sucrose water containing AMR or probiotic bacteria.

Fresh faecal samples were collected on specified days (**Fig. 5D**). Approximately 100 mg of faeces per mouse was suspended in 2 mL of normal saline and plated onto CHROMagar™ supplemented with antibiotics to detect bacteria carrying AMR plasmids (on CTX+ VAN plates) or the probiotic plasmid (on TET/FOS = VAN plates). Vancomycin (5 µg/mL) was used in agar plates to suppress Gram-positive Enterococcus growth while testing mouse faecal samples and this concentration of VAN does not affect Gram-negative *E. coli* growth. Following 18 hours of incubation at 37 °C, bacterial counts were expressed as colony-forming units per 100 mg of faeces. Isolates were further analysed by PCR to confirm plasmid-associated markers, including replicon types and AMR genes, as previously described [33].

After 96 hours of co-housing, both probiotic-treated (Gr10–12) and untreated control mice (Gr7–9) were challenged orally with cefotaxime (CTX) to select for AMR bacteria in the gut (**Fig. 5D**).

### The probiotic plasmid pPB1.1 protects the murine gut microbiota from invasion by diverse AMR plasmids

To assess the capacity of pPB1.1 to limit horizontal acquisition of AMR plasmids *in vivo*, mice treated with pPB1.1 were co-housed with mice carrying conjugative AMR plasmids representing three major incompatibility groups. Untreated mice served as controls, and all animals were maintained in groups of three as biological replicates. Testing of mouse gut microbiota before colonisation with AMR and probiotic plasmids revealed the presence of *E. coli* populations sensitive to all tested antibiotics (CTX, AZ, RIF, FOF, AMP, TET), providing a baseline for monitoring plasmid acquisition dynamics.

Mice were efficiently colonised by AMR and probiotic plasmids, as assigned by experimental group, provided in sucrose water made available *ad libitum* for 3 nights (**Fig. 6A, B)**. Both AMR and probiotic plasmids were efficiently established in the gut following administration in sucrose water (**Fig. 6A, B**) and were subsequently detected in the intestinal *E. coli* population, suggesting the self-transferable nature of both AMR and probiotic plasmids *in vivo*. *E. coli* NH78Rf carrying AMR plasmids were provided again for 2 nights (D9, 10) to AMR colonised mice, to maintain AMR plasmid colonisation in the mouse gut. J53Az carrying pPB1.1 was provided to probiotic-treated mice every night from D9 to D13 to maintain probiotic plasmid colonisation.

**Fig. 6.**
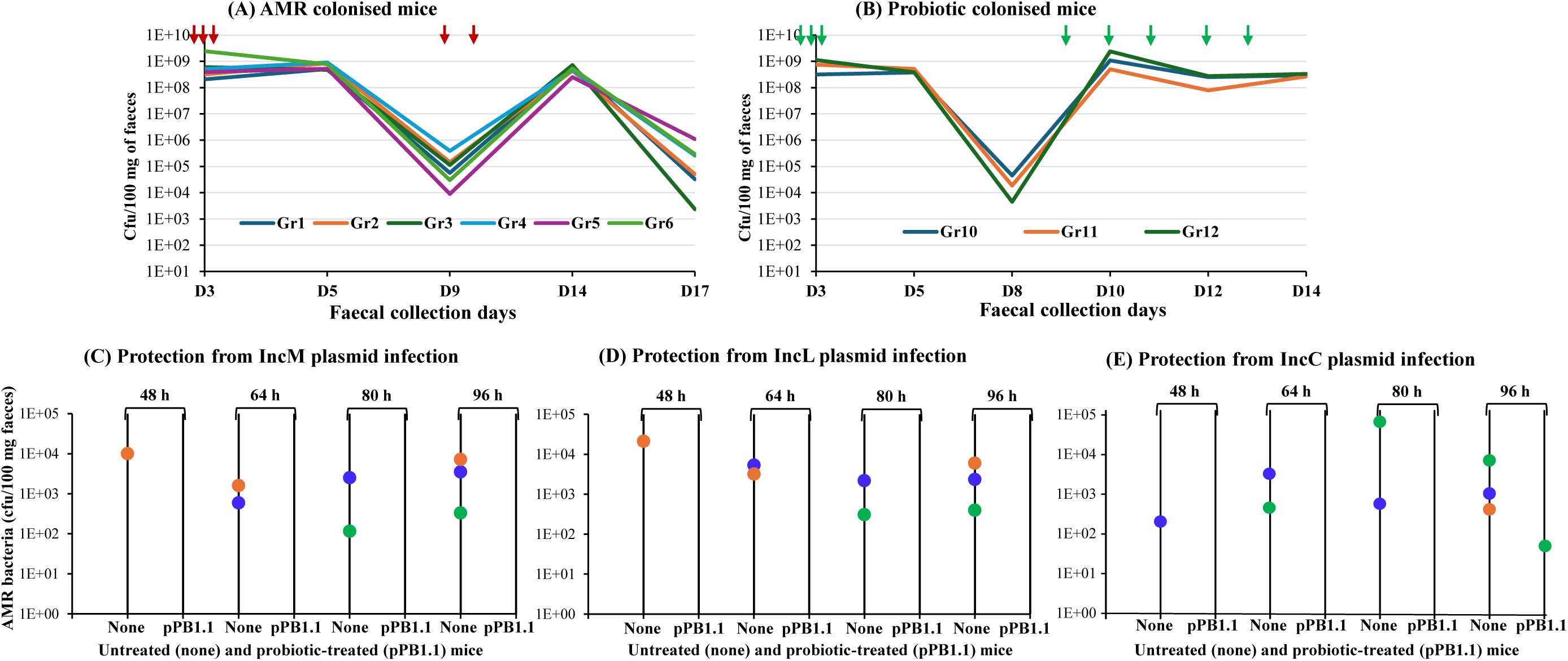
Probiotic plasmid protects the mouse gut from the acquisition of AMR plasmids. Six groups of mice were colonised with *E. coli* carrying AMR plasmids, and three groups of mice were colonised with *E. coli* carrying a probiotic plasmid (pPB1.1). Vertical red and green arrows represent the doses of AMR and probiotic plasmids, respectively. AMR and probiotic bacterial counts in AMR colonised (**A**, groups 1-6) and probiotic colonised (**B**, groups 10-12) mice, during the experiment. Data represent the average cfu values from three mice in each group. (**C-E**) AMR-colonised mice were co-housed with untreated mice (none) or probiotic-treated mice (pPB1.1). AMR plasmids were detected in mice from the untreated groups (none) after different co-housing times. Three different coloured circles represent three different mice in each group; blue, orange, and green filled circles represent mouse-1, mouse-2 and mouse-3, respectively, in each group.

Fresh faeces were collected and examined on specific antibiotic-containing CHROMagar^TM^ plates (VAN 5 µg/mL with CTX at 8 µg/mL for AMR plasmids and FOF at 200 µg/mL or TET at 10 µg/mL for probiotic plasmid) to measure colonisation, with serial dilution used to quantify.

In untreated control mice, AMR plasmids were rapidly acquired during co-housing, with detectable transfer within 48 h, and all untreated-control mice became AMR-infected within 96 h of exposure through coprophagy (**Fig. 6C-E**). AMR plasmids have been detected in the original NH78Rf strain (AMR-delivery bacteria) and in mouse gut *E. coli* populations. This suggests that the untreated mouse acquired AMR plasmids from the infected mouse during co-housing, and AMR plasmids then subsequently spread to the mouse gut *E. coli* and established well in the mouse gut environment. In contrast, only a single animal acquired an IncC plasmid at very low levels after prolonged exposure, and this plasmid was confined to the donor strain (*E. coli* NH78Rf), without detectable spread into resident or recipient *E. coli* populations, and all other probiotic-treated mice were completely protected from the acquisition of AMR plasmids (**Fig. 6C-E**).

PCR analysis of faecal samples confirmed the presence of AMR plasmid DNA in all animals at 96 h, indicating equivalent exposure across experimental groups. However, the absence of culturable AMR bacteria in probiotic-treated mice demonstrates that pPB1.1 does not prevent ingestion but instead blocks successful plasmid establishment and propagation. Consistent with this, pPB1.1 was observed to transfer into endogenous *E. coli* populations in the absence of antibiotic selection, becoming dominant among susceptible strains. This suggests that pPB1.1 efficiently occupies the available bacterial niche within the gut microbiota and may mediate the specific exclusion of related plasmids from entering.

Importantly, this protective effect was maintained under antibiotic selection. As a challenge of protection, both probiotic-treated (AMR culture-negative but PCR-positive) and untreated control mice (AMR culture-positive after exposure) were provided with cefotaxime (CTX) antibiotic in drinking sucrose water for 48 h to select for any mouse gut bacterial populations carrying AMR plasmids. Following cefotaxime exposure, high levels of CTX-resistant bacteria emerged in untreated control mice, whereas only sporadic and very low-level resistant populations were detected in a minority of pPB1.1-treated animals (**Fig. 7**). This indicates that pPB1.1 not only prevents initial plasmid acquisition but also substantially restricts the expansion of AMR-bearing populations under selective pressure.

**Fig. 7.**
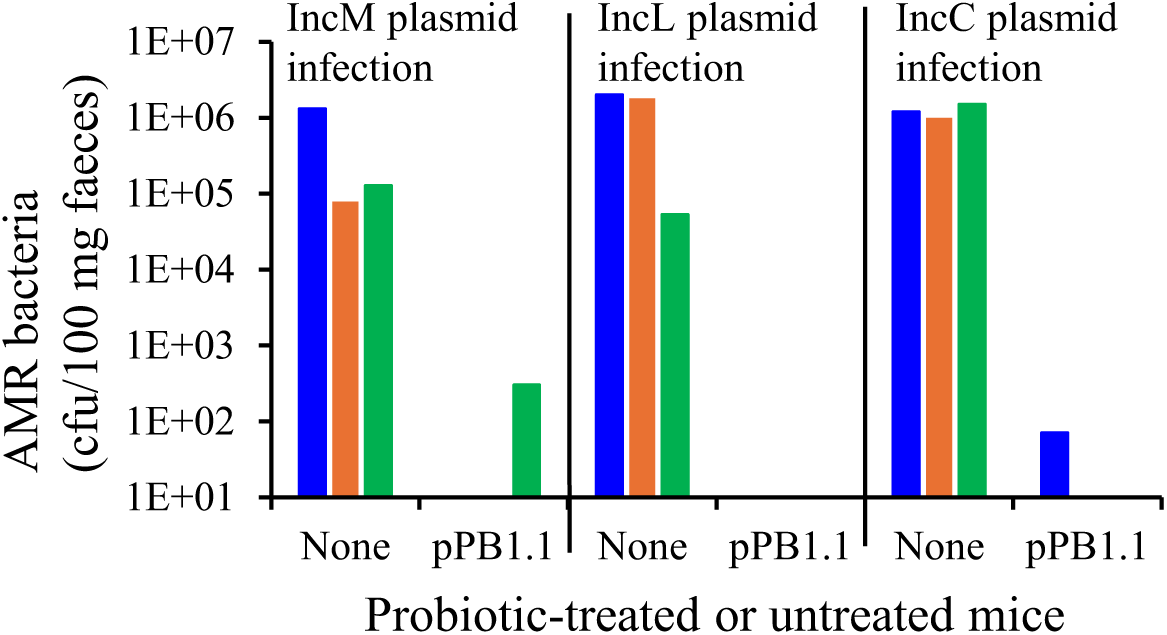
Test of protection after challenge with two doses of the cefotaxime (CTX) antibiotic. Probiotic plasmid-treated (pPB1.1) and untreated (None) mice were provided with CTX antibiotic in sucrose water for 2 days after 96 h co-housing with AMR plasmid colonised mice. After that, mouse faeces were collected, and CTX-resistant bacteria were counted from the CHROM Agar-CTX plates. Different colours represent different mice in each group; blue, orange, and green bars represent mouse-1, mouse-2, and mouse-3, respectively, in each group.

### Protection from AMR plasmid acquisition is independent of the delivery strain and plasmid backbone

To determine whether the observed microbiota protection effect of pPB1.1 was attributable to the delivery host (*E. coli* J53Az) or the IncM plasmid backbone (pJIMK46) used in its construction, we performed control co-housing experiments in the absence of pPB1.1-specific exclusion determinants. Mice colonised with IncM AMR plasmid were co-housed with mice colonised with J53Az alone, and mice colonised with IncL or IncC AMR plasmids were co-housed with J53Az carrying the backbone plasmid pJIMK46 (Table 1, which only has the IncM exclusion gene and no exclusion genes for IncC or IncL plasmids).

In contrast to the strong protection conferred by pPB1.1, AMR plasmid transfer occurred rapidly in all control groups. AMR-positive bacteria were detected in all mice within 48 h of co-housing (**Fig. 8**), demonstrating efficient horizontal plasmid transmission irrespective of the presence of J53Az or the IncM backbone plasmid alone. These findings indicate that neither the delivery strain nor the plasmid backbone provides measurable protection to AMR plasmid acquisition under these conditions.

**Fig. 8.**
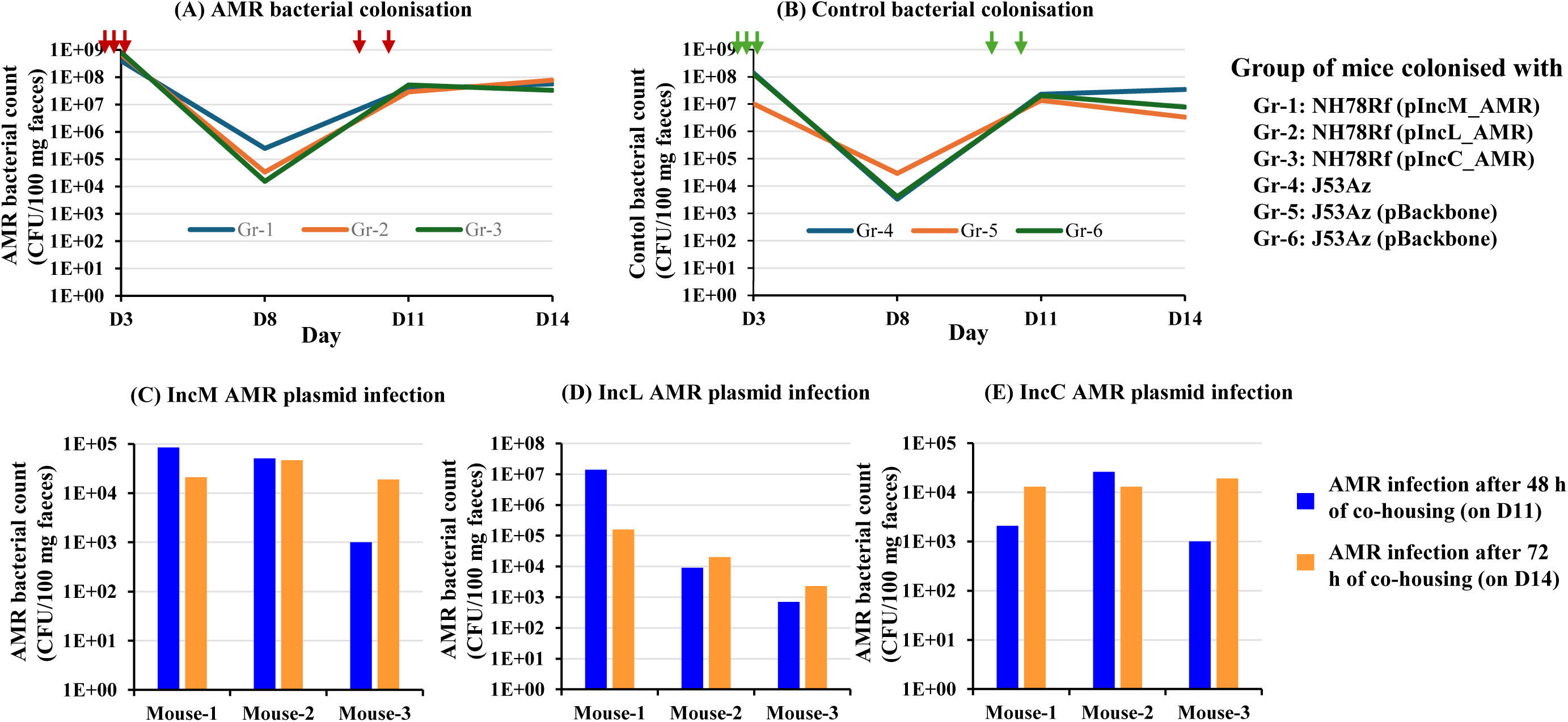
Effects of the probiotic plasmid delivery bacteria and the backbone plasmid in the acquisition of AMR plasmids during co-housing. **(A)** Groups of mice colonised with three different AMR plasmid carrying bacteria. (**B**) Groups of mice colonised with either probiotic plasmid delivery bacteria (Gr-4) or delivery bacteria carrying the backbone plasmid (pJIMK46) used to construct the probiotic plasmid pPB1.1 (Gr-5, 6). Vertical arrows represent the AMR and control bacterial doses (A, B). (**C**) IncM AMR plasmid acquisition in the mouse group (Gr-4), after co-housing with IncM AMR plasmid colonised mice (Gr-1). The mouse group-4 was colonised with the probiotic delivery bacteria J53Az. (**D, E**) IncL and IncC AMR plasmid acquisition in the mouse groups (Gr-5 and 6, respectively), after co-housing with IncL or IncC AMR plasmid colonised mice (Gr-2 and 3, respectively).

Collectively, these results demonstrate that the protective phenotype observed with pPB1.1 is specifically mediated by its engineered genetic components, plasmid exclusion mechanisms, rather than by nonspecific effects of the host strain or vector backbone. This reinforces the conclusion that targeted plasmid-based exclusion systems can confer robust and significant protection from the spread of AMR plasmids within complex microbial communities.

## Discussion

Our approach and supporting data illustrate a completely new strategy to exclude conjugative plasmids from bacterial ecosystems such as the mammalian gut and prevent the acquisition and spread of AMR genes, providing active protection against *de novo* acquisition and a perfect partner to plasmid curing strategies [28, 33].

Our prototype ‘protective’ plasmid has a high-efficiency conjugative backbone carrying *exc* from several clinically significant AMR plasmid families (IncM, IncL, and IncC), effectively shielding the mouse gut microbiota and preventing the acquisition of diverse target AMR plasmids.

This proof-of-principle study focused on IncM, IncL, and IncC plasmids, as three of the most important plasmid types driving modern multidrug resistance in bacteria [5]. They are relatively broad-host-range and well-characterised. Among others, IncM plasmids carry *bla*_IMP-4_, *bla*_NDM_ carbapenemase-encoding genes and *bla*_CTX-M_ extended-spectrum β-lactamase (ESBL)-encoding genes; IncL is known to disseminate the *bla*_OXA-48_ carbapenemase gene; IncC carries *bla*_NDM_ and *bla*_IMP-4_ carbapenemase genes, *bla*_CTX_ ESBL, and *bla*_CMY-2_ Amp-C β-lactamase genes [5, 7, 45]. While IncM and IncL facilitate resistance transfer among Enterobacteriaceae, IncC are also found in *Vibrio* and *Yersinia* species [46–49]. All of these factors prompted us to develop a preventive probiotic plasmid that targets these broad-host-range vehicles, providing a comprehensive shield against the dissemination of critical ESBL and carbapenemase genes.

To harness plasmid exclusion systems, our first step was to identify the genomic epidemiology and functional relatedness of exclusion systems in the target plasmids. Entry exclusion for the IncC plasmid is very well characterised and is known to inhibit the transfer of closely related IncA plasmid types [13]. All IncC plasmids in GenBank have a single Exc variant, IncA_Exc differs from IncC_Exc but is closely related (Supplementary **Fig. S1**). Exclusion genes for IncM and IncL plasmids are annotated in the plasmid sequences, but their function in plasmid exclusion has not been characterised. Here, we experimentally showed that the predicted Exc_M and Exc_L from the cloned mini-exclusion plasmids strongly inhibit the entry of their respective plasmids, confirming their exclusion properties (**Fig. 1**). Both Exc_M and Exc_L have four and five variants, respectively, with very minor amino acid changes. We demonstrated that one variant of Exc_M and Exc_L can inhibit the conjugation transfer of the IncM and IncL plasmids with the most distant Exc variant, confirming that these variations do not significantly reduce the exclusion of any IncM and IncL plasmids with Exc variants. That made it easy to choose a single Exc variant from each plasmid type for constructing our preventive plasmid.

The next challenge was to synthesise and clone multiple *exc* genes with their native promoter and RBS into a conjugation-efficient plasmid backbone, while ensuring proper expression of the *exc* genes from the new construct and that it does not impose additional fitness costs on bacteria.

To achieve this, we commercially synthesised the gBlock of *excC*-*excL*-*fosA3* with their native promoter and used it as a template for high-fidelity amplification using Phusion DNA polymerase, followed by insertion into the backbone of a high-conjugation-efficiency IncM plasmid derivative.

Whole-genome sequencing of the resulting probiotic plasmid, pPB1.1, identified a single nucleotide deletion in the promoter spacer region of the *exc_C* gene. While the exact sequence of the promoter region may vary, the spacer length between −10 and −35 hexamer elements is a critical determinant of promoter strength and transcriptional efficiency [50–51]. The spacer length of the *excC* putative promoter in the IncC plasmid is unusually long at 22 bp. A deletion of an A nucleotide in the spacer, reducing the length to 21 bp in pPB1.1, may contribute to the relatively high-level expression of the *excC* gene from pPB1.1 compared to that of the native IncC plasmid (**Fig. 3C**).

Similarly, *exc_L* also exhibited higher expression in pPB1.1 compared to its native IncL plasmid. In their original plasmid environments (IncC and IncL), *exc* gene expression is likely modulated by plasmid-encoded regulatory proteins. However, these regulatory elements were not included in pPB1.1, which is based on an IncM backbone. The absence of such regulators may permit constitutive expression of the *exc* genes, further contributing to their elevated expression in the engineered construct (**Fig. 3B**).

Importantly, despite the increased expression of both *excC* and *excL*, no significant additional fitness cost was observed in *E. coli* carrying pPB1.1 (**Fig. 3D**).

For the development of probiotic exclusion plasmids targeting other clinically important antimicrobial resistance (AMR) plasmids, it is essential to thoroughly characterise the corresponding entry exclusion systems, including their exclusion efficiency against target plasmids and the optimal constitutive expression levels of the exclusion genes. Achieving effective plasmid exclusion requires careful optimisation of exclusion gene expression within the construct. If the native promoter does not provide sufficient expression, exclusion genes should be evaluated under a range of moderate to strong constitutive promoters naturally associated with these plasmids. We have previously identified and characterised a promoter strength hierarchy suitable for this purpose, ranked as PcS > ISEcp1 Pout > PcWTGN-10 > Ptac [52]. Such optimisation is expected to maximise exclusion activity while minimising potential fitness costs to the host strain.

The broad host range and efficient *in vivo* conjugation of the probiotic plasmid are critical for its successful dissemination throughout the gut microbiota and for achieving robust protection against the acquisition of AMR plasmids. Previously, we identified a conjugative plasmid capable of efficient *in vivo* transfer across multiple bacterial species, which we used to construct AMR-curing plasmids that effectively eliminate AMR *in vivo* [33]. Building on this, we utilised the IncM plasmid backbone to develop the probiotic plasmid PB1.1. This construct provides complete protection against the acquisition of four major AMR plasmid types, IncL, IncM, IncA, and IncC, as demonstrated *in vitro* liquid mating assays (**Fig. 4**).

To evaluate its protective potential *in vivo*, we established a mouse model in which AMR-infected mice were co-housed with either probiotic-treated or untreated mice. This setup allowed us to assess whether probiotic treatment could protect against AMR transmission via incidental exposure to contaminated faeces through coprophagy. As expected, probiotic-treated mice exhibited complete protection against AMR colonisation, in contrast to untreated controls. Within 96 hours of co-housing (∼8 hours per day), all mice in untreated groups were colonised with bacteria carrying the tested AMR plasmid types (IncC, IncL, and IncM).

PCR analysis of faecal samples from probiotic-treated mice detected AMR plasmid genes after 96 hours; however, no corresponding AMR-carrying bacteria were recoverable on selective agar plates. This suggests that the probiotic plasmid had already disseminated widely across the gut microbiota, thereby preventing stable establishment and spread of incoming AMR plasmids. Indeed, we observed extensive spread of PB1.1 following repeated dosing.

To further test the durability of protection, all mice were challenged with cefotaxime after ∼96 hours of co-housing to reveal any undetected AMR populations. Following antibiotic exposure, untreated mice exhibited high levels of AMR bacterial colonisation (∼10⁶ CFU/mg of faeces, **Fig. 7**). In contrast, only two out of nine probiotic-treated mice showed minimal colonisation (∼10² CFU/mg), confirming the strong protective effect of pPB1.1.

The observed protection conferred by the probiotic plasmid pPB1.1 could potentially be attributed either to colonisation by the delivery *E. coli* strain or to genetic features of the plasmid backbone independent of the exclusion genes. However, we ruled out these possibilities by demonstrating that mice colonised with the delivery strain alone, or with the strain carrying only the backbone plasmid used to construct pPB1.1, showed no protection against AMR plasmid acquisition (**Fig. 8**).

Notably, the entire study was conducted without antibiotic selection to support colonisation or plasmid dissemination, highlighting the intrinsic stability and transfer efficiency of PB1.1. Although the current construct contains antibiotic resistance markers (*fosA3* and *tetA*) for experimental selection, future therapeutic versions will be antibiotic-sensitive, as our findings demonstrate that neither colonisation nor plasmid spread requires antibiotic pressure.

The plasmid stability system of pPB1.1, based on a toxin-antitoxin (TA) module, was intentionally modified by deleting the toxin gene. This engineering renders the plasmid unstable in the absence of selective pressure. Consistent with our previous findings, the *pemK*-deleted version of the IncM backbone is rapidly lost from bacterial populations within one week after cessation of probiotic administration [33]. In the present study, we similarly observed that pPB1.1 titers declined significantly within three days of stopping ingestion (**Fig. 6B**).

Because pPB1.1 is derived from a natural conjugative plasmid, there is a theoretical risk of recombination with endogenous AMR plasmids. However, we did not detect any recombination events in this study or in our previous AMR-curing experiments using interference plasmids under antibiotic selection [33]. To further mitigate this risk, all transposable elements were deliberately removed during plasmid construction. In addition, no antibiotics were used during the experimental protocol, thereby avoiding selective pressures that could promote recombination.

Leveraging natural plasmid exclusion systems to block the acquisition and spread of undesirable plasmids represents both a novel and potentially transformative strategy. This approach could be particularly valuable for individuals at high risk of exposure to AMR, such as patients in healthcare settings or travellers to regions with high prevalence of resistant pathogens.

In this context, the probiotic plasmid functions analogously to a “vaccine” against AMR, virulence determinants, and other undesirable self-transmissible plasmids. Unlike conventional interventions, current AMR control strategies, including antibiotic stewardship, surveillance, isolation, and antibiotic development, have remained largely unchanged for decades. Our approach introduces a fundamentally new paradigm by preventing initial plasmid invasion, thereby interrupting the cycle of AMR transmission. This strategy could enable clinicians to safely use existing antibiotics while preserving the efficacy of newly developed agents. Ultimately, it offers a dual benefit: limiting the spread of resistance and protecting vulnerable patients from AMR-associated morbidity and mortality.

Collectively, these findings demonstrate that a conjugative probiotic plasmid can disseminate within the gut microbiota and confer robust, population-level resistance to invasion by diverse AMR plasmids. The ability of pPB1.1 to combine efficient horizontal transfer with broad exclusion activity highlights its potential as a scalable strategy to limit the spread of antimicrobial resistance in complex microbial ecosystems.

## Acknowledgements

We thank Ruth Hall for providing plasmid RA1. Graphical abstract and Fig. 2 have been made using BioRender.

## Conflict of interest

We declare conflict of interest. The data provided in this manuscript are used in a patent application (WO2023092187). MK and JI are the inventors of that patent application.

## Funding

The work was supported by the Australian National Health and Medical Research Council (NHMRC) project grant GNT1145914 to MK and JRI, and Investigator grant GNT1197534 to JRI.

## Data availability

Data generated and presented in the manuscript has been found in the manuscript and in the supplementary information file.

## Authors contributions

M.K. conceived the study, constructed probiotic plasmids, performed *in vitro* and *in vivo* experiments, conducted data analysis, provided supervision and overall guidance, secured funding and was responsible for writing the original draft, review & editing.

A.Y.W. and J.J. contributed to the animal experiments in testing probiotic plasmid in mouse.

J.R.I. provided study resources, secured funding, performed project administration, supervised and contributed to review & editing.

## Notes

### Competing Interest Statement

We declare competing interest. The data provided in this manuscript are used in a patent application (WO2023092187). MK and JI are the inventors of that patent application.

## References

1. Antimicrobial Resistance Collaborators, C. Global burden of bacterial antimicrobial resistance in 2019: a systematic analysis. Lancet 2022, 399 (10325), 629–655.

2. GBD 2021 Antimicrobial Resistance Collaborators, C. Global burden of bacterial antimicrobial resistance 1990-2021: a systematic analysis with forecasts to 2050. Lancet 2024, 404 (10459), 1199–1226.

3. O’Neill, J. Antimicrobial Resistance: Tackling a crisis for the health and wealth of nations. London: *Review on Antimicrobial resistance*; 2014.

4. Shallcross, L.J.; Davies, S.C. The World Health Assembly resolution on antimicrobial resistance. J Antimicrob Chemother 2014, 69 (11), 2883–5.

5. Carattoli, A. Resistance plasmid families in *Enterobacteriaceae*. Antimicrob Agents Chemother 2009, 53 (6), 2227–38.

6. Liu, Y.Y.; Wang, Y.; Walsh, T.R.; Yi, L.X.; Zhang, R.; Spencer, J.; Doi, Y.; Tian, G.; Dong, B.; Huang, X.; Yu, L.F.; Gu, D.; Ren, H.; Chen, X.; Lv, L.; He, D.; Zhou, H.; Liang, Z.; Liu, J.H.; Shen, J. Emergence of plasmid-mediated colistin resistance mechanism MCR-1 in animals and human beings in China: a microbiological and molecular biological study. Lancet Infect Dis 2016, 16 (2), 161–8.

7. Partridge, S.R.; Kwong, S.M.; Firth, N.; Jensen, S.O. Mobile genetic elements associated with antimicrobial resistance. Clin Microbiol Rev 2018, 31 (4), e00088–17.

8. Pilla, G.; Tang, C.M. Going around in circles: virulence plasmids in enteric pathogens. Nat Rev Microbiol 2018, 16 (8), 484–495.

9. Smillie, C.; Garcillan-Barcia, M.P.; Francia, M.V.; Rocha, E.P.; de la Cruz, F. Mobility of plasmids. Microbiol Mol Biol Rev 2010, 74 (3), 434–52.

10. Hayes, F. Toxins-antitoxins: plasmid maintenance, programmed cell death, and cell cycle arrest. Science 2003, 301 (5639), 1496–99.

11. Kamruzzaman, M.; Wu, A.Y.; Iredell, J.R. Biological functions of type II toxin-antitoxin systems in bacteria. Microorganisms 2021, 9 (6).

12. Garcillan-Barcia, M.P.; de la Cruz, F. Why is entry exclusion an essential feature of conjugative plasmids? Plasmid 2008, 60 (1), 1–18.

13. Humbert, M.; Huguet, K.T.; Coulombe, F.; Burrus, V. Entry exclusion of conjugative plasmids of the IncA, IncC, and related untyped incompatibility groups. J Bacteriol 2019, 201 (10), e00731–18.

14. Kamruzzaman, M.; Mathers, A.J.; Iredell, J.R. A novel plasmid entry exclusion system in pKPC_UVA01, a promiscuous conjugative plasmid carrying the *bla*_KPC_ carbapenemase gene. Antimicrob Agents Chemother 2022, 66 (3), e0232221.

15. Carattoli, A.; Bertini, A.; Villa, L.; Falbo, V.; Hopkins, K.L.; Threlfall, E.J. Identification of plasmids by PCR-based replicon typing. J Microbiol Methods 2005, 63 (3), 219–28.

16. Novick, R.P. Plasmid incompatibility. Microbiol Rev 1987, 51 (4), 381–95.

17. Audette, G.F.; Manchak, J.; Beatty, P.; Klimke, W.A.; Frost, L.S. Entry exclusion in F-like plasmids requires intact TraG in the donor that recognizes its cognate TraS in the recipient. Microbiology (Reading) 2007, 153 (Pt 2), 442–51.

18. Gunton, J.E.; Ussher, J.E.; Rooker, M.M.; Wetsch, N.M.; Alonso, G.; Taylor, D.E. Entry exclusion in the IncHI1 plasmid R27 is mediated by EexA and EexB. Plasmid 2008, 59 (2), 86–101.

19. Haase, J.; Kalkum, M.; Lanka, E. TrbK, a small cytoplasmic membrane lipoprotein, functions in entry exclusion of the IncPα plasmid RP4. J Bacteriol 1996, 178 (23), 6720–9.

20. Hartskeerl, R.A.; Hoekstra, W.P. Exclusion in IncI-type *Escherichia coli* conjugations: the stage of conjugation at which exclusion operates. Antonie Van Leeuwenhoek 1984, 50 (2), 113–24.

21. Pohlman, R.F.; Genetti, H.D.; Winans, S.C. Entry exclusion of the IncN plasmid pKM101 is mediated by a single hydrophilic protein containing a lipid attachment motif. Plasmid 1994, 31 (2), 158–65.

22. Sakuma, T.; Tazumi, S.; Furuya, N.; Komano, T. ExcA proteins of IncI1 plasmid R64 and IncIgamma plasmid R621a recognize different segments of their cognate TraY proteins in entry exclusion. Plasmid 2013, 69 (2), 138–45.

23. Ardal, C.; Balasegaram, M.; Laxminarayan, R.; McAdams, D.; Outterson, K.; Rex, J.H.; Sumpradit, N. Antibiotic development - economic, regulatory and societal challenges. Nat Rev Microbiol 2020, 18 (5), 267–274.

24. Ventola, C.L. The antibiotic resistance crisis: part 1: causes and threats. P T 2015, 40 (4), 277–83.

25. Buckner, M.M.C.; Ciusa, M.L.; Piddock, L.J.V. Strategies to combat antimicrobial resistance: anti-plasmid and plasmid curing. FEMS Microbiol Rev 2018, 42 (6), 781–804.

26. Pursey, E.; Sunderhauf, D.; Gaze, W.H.; Westra, E.R.; van Houte, S. CRISPR-Cas antimicrobials: Challenges and future prospects. PLoS Pathog 2018, 14 (6), e1006990.

27. Rodrigues, M.; McBride, S.W.; Hullahalli, K.; Palmer, K.L.; Duerkop, B.A. Conjugative delivery of CRISPR-Cas9 for the selective depletion of antibiotic-resistant *Enterococci*. Antimicrob Agents Chemother 2019, 63 (11).

28. Wang, P.; He, D.; Li, B.; Guo, Y.; Wang, W.; Luo, X.; Zhao, X.; Wang, X. Eliminating mcr-1-harbouring plasmids in clinical isolates using the CRISPR/Cas9 system. J Antimicrob Chemother 2019, 74 (9), 2559–65.

29. Fernandez-Lopez, R.; Machon, C.; Longshaw, C.M.; Martin, S.; Molin, S.; Zechner, E.L.; Espinosa, M.; Lanka, E.; de la Cruz, F. Unsaturated fatty acids are inhibitors of bacterial conjugation. Microbiology (Reading) 2005, 151 (Pt 11), 3517–3526.

30. Getino, M.; Fernandez-Lopez, R.; Palencia-Gandara, C.; Campos-Gomez, J.; Sanchez-Lopez, J.M.; Martinez, M.; Fernandez, A.; de la Cruz, F. Tanzawaic acids, a chemically novel set of bacterial conjugation inhibitors. PLoS One 2016, 11 (1), e0148098.

31. Getino, M.; Sanabria-Rios, D.J.; Fernandez-Lopez, R.; Campos-Gomez, J.; Sanchez-Lopez, J.M.; Fernandez, A.; Carballeira, N.M.; de la Cruz, F. Synthetic fatty acids prevent plasmid-mediated horizontal gene transfer. mBio 2015, 6 (5), e01032–15.

32. Palencia-Gandara, C.; Getino, M.; Moyano, G.; Redondo, S.; Fernandez-Lopez, R.; Gonzalez-Zorn, B.; de la Cruz, F. Conjugation inhibitors effectively prevent plasmid transmission in natural environments. mBio 2021, 12 (4), e0127721.

33. Kamruzzaman, M.; Shoma, S.; Thomas, C.M.; Partridge, S.R.; Iredell, J.R. Plasmid interference for curing antibiotic resistance plasmids in vivo. PLoS One 2017, 12 (2), e0172913.

34. Matsumura, Y.; Peirano, G.; Pitout, J.D.D. Complete genome sequence of *Escherichia coli* J53, an azide-resistant laboratory strain used for conjugation experiments. Genome Announc 2018, 6 (21).

35. de la Cruz, F.; Grinsted, J. Genetic and molecular characterization of Tn*21*, a multiple resistance transposon from R100.1. Journal of bacteriology 1982, 151 (1), 222–28.

36. Espedido, B.A.; Partridge, S.R.; Iredell, J.R. *bla*_IMP-4_ in different genetic contexts in Enterobacteriaceae isolates from Australia. Antimicrob Agents Chemother 2008, 52 (8), 2984–7.

37. Papagiannitsis, C.C.; Kutilova, I.; Medvecky, M.; Hrabak, J.; Dolejska, M. Characterization of the complete nucleotide sequences of IncA/C(2) plasmids carrying In809-like integrons from Enterobacteriaceae isolates of wildlife origin. Antimicrob Agents Chemother 2017, 61 (9).

38. Agyekum, A.; Fajardo-Lubian, A.; Ai, X.; Ginn, A.N.; Zong, Z.; Guo, X.; Turnidge, J.; Partridge, S.R.; Iredell, J.R. Predictability of phenotype in relation to common β-lactam resistance mechanisms in *Escherichia coli* and *Klebsiella pneumoniae*. J Clin Microbiol 2016, 54 (5), 1243–50.

39. Fajardo-Lubian, A.; Ben Zakour, N.L.; Agyekum, A.; Qi, Q.; Iredell, J.R. Host adaptation and convergent evolution increases antibiotic resistance without loss of virulence in a major human pathogen. PLoS Pathog 2019, 15 (3), e1007218.

40. Murphy, K.C.; Campellone, K.G. Lambda red-mediated recombinogenic engineering of enterohemorrhagic and enteropathogenic *E. coli*. BMC Mol Biol 2003, 4, 11.

41. Qi, Q.; Kamruzzaman, M.; Iredell, J.R. The *higBA*-type toxin-antitoxin system in IncC plasmids is a mobilizable ciprofloxacin-inducible system. mSphere 2021, 6 (3), e0042421.

42. Bonnin, R.A.; Nordmann, P.; Carattoli, A.; Poirel, L. Comparative genomics of IncL/M-type plasmids: evolution by acquisition of resistance genes and insertion sequences. Antimicrob Agents Chemother 2013, 57 (1), 674–6.

43. Carattoli, A.; Seiffert, S.N.; Schwendener, S.; Perreten, V.; Endimiani, A. Differentiation of IncL and IncM plasmids associated with the spread of clinically relevant antimicrobial resistance. PLoS One 2015, 10 (5), e0123063.

44. Kamruzzaman, M.; Iredell, J. A ParDE-family toxin antitoxin system in major resistance plasmids of Enterobacteriaceae confers antibiotic and heat tolerance. Sci Rep 2019, 9 (1), 9872.

45. Carattoli, A. Plasmids and the spread of resistance. Int J Med Microbiol 2013, 303 (6-7), 298–304.

46. Carraro, N.; Rivard, N.; Ceccarelli, D.; Colwell, R.R.; Burrus, V. IncA/C conjugative plasmids mobilize a new family of multidrug resistance islands in clinical *Vibrio cholerae* non-O1/non-O139 isolates from Haiti. mBio 2016, 7 (4).

47. Galimand, M.; Guiyoule, A.; Gerbaud, G.; Rasoamanana, B.; Chanteau, S.; Carniel, E.; Courvalin, P. Multidrug resistance in *Yersinia pestis* mediated by a transferable plasmid. N Engl J Med 1997, 337 (10), 677–80.

48. Pan, J.C.; Ye, R.; Wang, H.Q.; Xiang, H.Q.; Zhang, W.; Yu, X.F.; Meng, D.M.; He, Z.S. *Vibrio cholerae* O139 multiple-drug resistance mediated by *Yersinia pestis* pIP1202-like conjugative plasmids. Antimicrob Agents Chemother 2008, 52 (11), 3829–36.

49. Weill, F.X.; Domman, D.; Njamkepo, E.; Tarr, C.; Rauzier, J.; Fawal, N.; Keddy, K.H.; Salje, H.; Moore, S.; Mukhopadhyay, A.K.; Bercion, R.; Luquero, F.J.; Ngandjio, A.; Dosso, M.; Monakhova, E.; Garin, B.; Bouchier, C.; Pazzani, C.; Mutreja, A.; Grunow, R.; Sidikou, F.; Bonte, L.; Breurec, S.; Damian, M.; Njanpop-Lafourcade, B.M.; Sapriel, G.; Page, A.L.; Hamze, M.; Henkens, M.; Chowdhury, G.; Mengel, M.; Koeck, J.L.; Fournier, J.M.; Dougan, G.; Grimont, P.A.D.; Parkhill, J.; Holt, K.E.; Piarroux, R.; Ramamurthy, T.; Quilici, M.L.; Thomson, N.R. Genomic history of the seventh pandemic of cholera in Africa. Science 2017, 358 (6364), 785–789.

50. Hawley, D.K.; McClure, W.R. Compilation and analysis of *Escherichia coli* promoter DNA sequences. Nucleic Acids Res 1983, 11 (8), 2237–55.

51. Jensen, P.R.; Hammer, K. The sequence of spacers between the consensus sequences modulates the strength of prokaryotic promoters. Appl Environ Microbiol 1998, 64 (1), 82–7.

52. Kamruzzaman, M.; Patterson, J.D.; Shoma, S.; Ginn, A.N.; Partridge, S.R.; Iredell, J.R. Relative strengths of promoters provided by common mobile genetic elements associated with resistance gene expression in Gram-negative bacteria. Antimicrob Agents Chemother 2015, 59 (8), 5088–91.

