## Supplementary table and figures for "Specific exclusion of conjugative plasmids from the gut microflora"

**Table S1. Primers used in this study**

| Primers | Sequences (5´-3´)* | Source |
|---|---|---|
| ExcL_F-XbaI | CG**TCTAGA**TCTTGGCACCGCGCTTAACT | This study |
| ExcL_R-HindIII | GC**AAGCTT**GCCTTTTCAGGCGGTTTGGA | This study |
| ExcM_F-XbaI | CG**TCTAGA**TGCGAGGATGCCGAACTATG | This study |
| ExcM_R-HindIII | GC**AAGCTT**CGCCATATTCGGCGGGTTA | This study |
| EXC1-F | <u>GAGCCTCTACGCCGACCTCATTCAACAGGTCCAGGGCGAC</u>GCTCTTGGCACCGCGCTTAA | This study |
| FOSA3-R | <u>TAGCCGGAAGTCGCCTTGACCCGCATGGCATAGGCCTAT</u>GCTGTGGATCTGCACGTTGAA | This study |
| PemK-tetA-F | <u>GGCTTGTCTCGCTTGACCCTACCGCAGGTCATGAGCAGCA</u>GAGCCTCTACGCCGACCTCA | This study |
| PemK-tetA_R | <u>TTCGAGTCGTTTGCCGCCGCGGGCTTTCATATCGATCGTC</u>GTAGCCGGAAGTCGCCTTGA | This study |
| ExcM_F_RT | AAGGACTTCAGGAGCCGCAAGC | This study |
| ExcM_R_RT | CCGGGCCAGGTCATAGTGCTC | This study |
| ExcL_F_RT | CTGTCACGCCTGTCTGTAGCCA | This study |
| ExcL_R_RT | GCAACAAACCACGCCAATGCC | This study |
| ExcC_F_RT | CTTCGGCGTTTGGGGTGCTC | This study |
| ExcC_R_RT | AGAGCCCCGACAACCAAAGCA | This study |

*Restriction sites are shown in bold, overlapping sequences with the insertion sites are underlined.

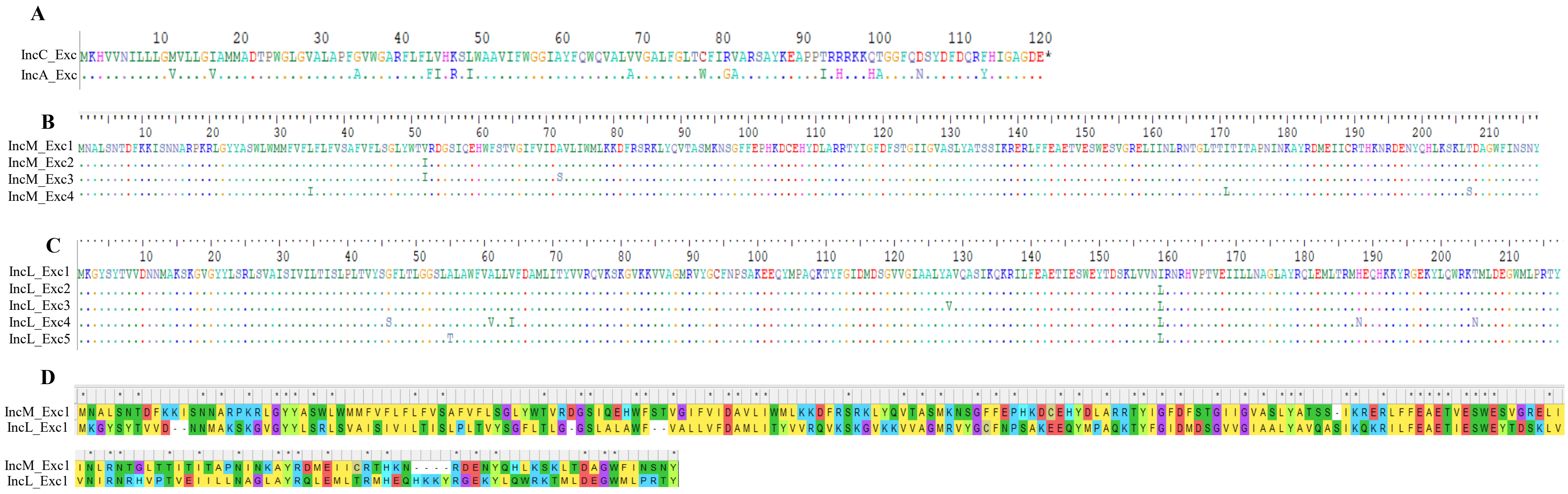

**Fig. S1. Amino acid sequence alignments of Exc proteins and their variants.** (**A**) IncC and IncA Exc, (**B**) IncM plasmid Exc variants, (**C**) IncL plasmid Exc variants, and (**D**) major Exc of IncM and IncL plasmids. Identical amino acids are shown by dots (A-C). Identical amino acids between Exc M and Exc L are shown by asterisks (D). Amino acid positions are shown by the numbers on top (A-C).

**A**

| | |
|---|---|
|  | -35 -10 |
| IncL_exc1 | **TTGCTG**CATTTGGTCGTGGAGCT**AAAAAT** |
| IncL_exc2 | ----------------------------- |
| IncL_exc3 | ----------------------------- |
| IncL_exc4 | ----------------------------- |
| IncL_exc5 | ----------------------------- |

**B**

| | |
|---|---|
|  | -35 -10 |
| IncM_exc1 | **TTGCTG**CGTTCGGTCGTGGAGCC**AAAAAT** |
| IncM_exc2 | ----------------G------------ |
| IncM_exc3 | ----------------------------- |
| IncM_exc4 | ----------------A---A-------- |
| IncL_excs | -------A--T-----------T------ |

**Fig. S2.** Alignment of predicted promoter sequences associated with the *exc* variants of IncL (**A**) and IncM and IncL plasmids (**B**). -35 and -10 sequences are shown in bold. Identical nucleotide sequences are shown by dashes.

**A. Synthesized e*xc4_M* with XbaI and HindIII sites**

CGTCTAGATGCGAGGATGCCGAACTATGTTATAAACTGGATTGGACAGAAGATGAACGATTCAGTGTTGGGTGATATGCAGAACCATGTT
CATGACATCTTTGCTGCGTTCGGTCGAGGAACCAAAAATATGAACCGGAACAACCCTAAACAATTTAATCCTGACGCAGGTGTAGATAAG
AGCAAAGATGGCATCAAAGGAGCTTAAATATGAATGCGTTATCAAACACTGACTTTAAGAAAATTAGTAACAATGCGCGTCCTAAAAGGC
TTGGCTACTACGCATCATGGCTCTGGATGATGTTTGTATTTATCTTCCTGTTCGTTAGCGCTTTCGTGTTTCTGAGTGGCTTATACTGGA
CGGTTAGAGATGGTTCAATCCAGGAGCACTGGTTTAGTACAGTCGGTATCTTTGTGATAGATGCCGTGCTGATCTGGATGTTGAAAAAGG
ACTTCAGGAGCCGCAAGCTCTACCAGGTTACGGCTTCAATGAAAAACTCTGGCTTCTTTGAGCCACATAAGGACTGCGAGCACTATGACC
TGGCCCGGAGGACTTACATAGGCTTTGACTTCAGTACTGGAATAATAGGAGTCGCATCCCTGTATGCAACATCCAGTATTAAACGTGAGC
GGCTATTTTTTGAAGCCGAAACAGTTGAATCATGGGAGTCAGTTGGCAGGGAGTTAATCATAAACCTGCGAAATACGGGGCTCACGACGC
TCACGATCACTGCTCCAAATATCAATAAAGCGTACAGGGATATGGAAATCATCTGCAGAACGCATAAAAATAGGGATGAAAACTATCAGC
ATCTGAAGAGTAAACTGTCAGACGCAGGATGGTTCATAAACAGCAATTATTGAGTTTAACCCGCCGAATATGGCGAAGCTTCG

**B. Synthesized e*xc4_L* with XbaI and HindIII sites**

CGTCTAGATCTTGGCACCGCGCTTAACTATATTTTCAAAACAGCAATGGAAAATGTACAAGCAGATTCAACAACTGGCCTTTGGTCAATG
ATAGGTATCCTGTTTGTCTATGCAAGGATGTGTACAGGTATGGTTGCTAGGGTATTTGCATTACCTGCTAGGATGCCTAACTACGTCATC
AGCTGGATTGGTAACAAGCATAACGATTCAATTCTTGGTGATATGCAGAATCATGTTCACGACCTGTTTGCTGCATTTGGTCGTGGAGCT
AAAAATACAGGTGGTCGTCCTTCTGGGCCGAAGAACTTCAACCCAACCAATACGGTTGATAAAGACAAAGATGGTATAGCGGGGGCGTGA
TATGAAAGGATATTCGTATACTGTTGTCGATAACAACATGGCTAAATCGAAAGGTGTTGGCTACTATCTGTCACGCCTGTCTGTTGCCAT
AAGTATAGTTATTTTAACAATAAGTCTTCCGTTGACTGTATATTCTAGCTTTCTTACTTTAGGTGGAAGCCTGGCATTGGCGTGGTTTGT
TGTTTTACTTATTTTCGATGCCATGCTGATCACATATGTTGTACGCCAGGTAAAAAGTAAAGGAGTCAAAAAGGTCGTTGCAGGCATGAG
AGTTTATGGATGCTTTAATCCATCGGCTAAAGAGGAACAATACATGCCGGCTCAGAAAACATATTTTGGTATAGACATGGACTCTGGTGT
TGTTGGGATTGCTGCCCTTTATGCTGTACAGGCCTCTATTAAGCAAAAGAGAATTTTGTTTGAGGCCGAGACAATAGAGTCTTGGGAGTA
TACTGATAGTAAACTTGTCGTGAATCTCCGGAACCGACATGTACCTACTGTTGAAATTATCTTGCTCAATGCCGGCCTAGCATATCGCCA
GCTGGAAATGTTAACCAGGATGAATGAGCAACATAAAAAATACAGGGGAGAAAAATATCTCCAGTGGCGAAAAAACATGCTTGATGAAGG
ATGGATGCTGCCACGAACGTATTGATGTTTACTTCCAAACCGCCTGAAAAGGCAAGCTTCG

**Fig. S3. Sequences of the synthesised *exc* gene variants for *exc4_M* (A) and *exc4_L* (B).** Underlined sequences are restriction enzyme cutting sites.

A

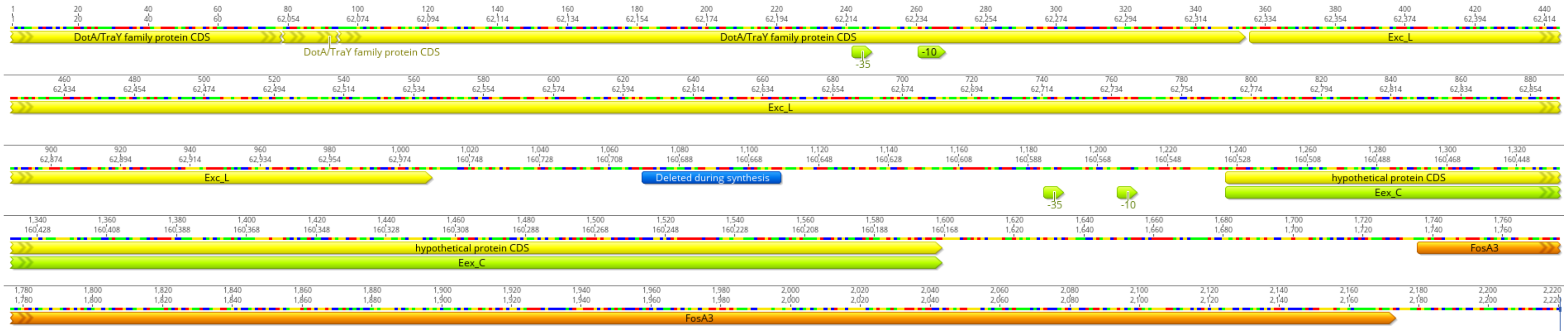

DotA/TraY family protein CDS
DotA/TraY family protein CDS
DotA/TraY family protein CDS
Exc_L
-35
-10
Exc_L
Exc_L
Deleted during synthesis
hypothetical protein CDS
Eex_C
-35
-10
hypothetical protein CDS
Eex_C
FosA3
FosA3

**B**

1 GCTCTTGGCACCGCGCTTAACTATATTTTCAAAACAGCAATGGAAAATGTACAGGCAGAT

61 TCAACAACTGGCCTTTGGTCAATGATAGGTATCCTTTTTGTCTATGCAAGGATGTGTACA

121 GGTATGGTCGCTAGGGTATTTGCATTACCTGCAAGGATGCCTAACTACGTCATCAGCTGG

181 ATTGGTAACAAGCATAACGATTCAATTCTTGGTGATATGCAGAATCATGTTCACGACCTG

241 TTTGCTGCATTTGGTCGTGGAGCTAAAAATACAGGTGGACGTCCTTCTGGGCCGAAGAAC

301 TTCAACCCAACCAATACGGTTGATAAAGACAAAGATGGTATAGCGGGGGCGTGATATGAA

361 AGGATATTCGTATACTGTTGTCGATAACAACATGGCTAAATCAAAAGGTGTTGGCTACTA

421 TCTGTCACGCCTGTCTGTAGCCATAAGTATAGTTATTTTAACAATAAGTCTTCCGTTGAC

481 TGTATATTCTGGCTTTCTTACTTTAGGTGGAAGCCTGGCATTGGCGTGGTTTGTTGCTCT

541 ACTTGTTTTCGATGCCATGCTGATCACATATGTTGTACGCCAGGTAAAAAGTAAGGGAGT

601 CAAAAAGGTCGTTGCAGGCATGAGAGTTTATGGATGCTTTAATCCATCGGCTAAAGAGGA

661 ACAATACATGCCGGCTCAGAAAACATATTTTGGTATAGATATGGACTCCGGTGTTGTTGG

721 TATTGCTGCCCTTTATGCTGTACAGGCCTCTATTAAGCAAAAGAGAATTTTGTTTGAGGC

781 CGAGACAATTGAGTCTTGGGAGTATACTGATAGTAAACTTGTCGTAAATATTCGGAACCG

841 ACATGTGCCTACTGTTGAAATAATCTTGCTCAATGCCGGCCTAGCATATCGCCAGCTGGA

901 AATGTTAACCAGGATGCATGAGCAACATAAAAAATACAGGGGTGAAAAATATCTCCAGTG

961 GCGAAAGACTATGCTTGATGAAGGTTGGATGCTACCACGAACGTATTGATGTTTTTCCAT

1021 AGTGTATGCGATTTGTTTGACATGTTCAACTATGTAATGCTTTGTCAATGAGGGCAAATT

1081 CTAATCGAAGAAGTTGCCAATGGAATGGTTGTTTAGCAAAAAATAGTCAAACTTTCACAG

1141 TTGCATTTTTTGATGCCGTGGTAAACTGGAGGTAAAGAGTAAGGGGGTTGATATGAAACA

1201 TGTGGTCAATATTCTTCTGCTGGGAATGGTGCTTCTGGGAATAGCCATGATGGCTGACAC

1261 ACCTTGGGGACTTGGTGTGGCGCTGGCTCCCTTCGGCGTTTGGGGTGCTCGTTTCCTATT

1321 TCTGGTTCACAAAAGCCTTTGGGCTGCTGTTATCTTTTGGGGAGGGATCGCTTACTTCCA

1381 GTGGCAAGTTGCTTTGGTTGTCGGGGCTCTGTTTGGACTAACGTGCTTTATCCGCGTTGC

1441 GCGGTCAGCATATAAAGAGGCACCACCTACCCGGCGCAGAAAAAAACAAACTGGTGGGTT

```
1501   TCAAGATTCCTATGATTTTGACCAGCGCTTCCATATTGGAGCTGGAGACGAATAAGGGGG

1561   CATAGAGCGGGGATTAGTGTGGCGAGGCGCAGGTTTCCGATGGCAGGCTAAACGCAAAAA

1621   ATGCGCTTTTTAGCCGGTGATGAGGTGAGGCCGGGAGGGGATGCGTCGCCGATCACAGTT

1681   TACAACAGGGTTTGATAACGGGAGGAAAAGTCATGCTGCAGGGATTGAATCATCTGACGC

1741   TGGCGGTCAGCGATCTGGCGTCAAGCCTGGCATTTTATCAGCAGTTACCTGGAATGCGCC

1801   TGCACGCCAGCTGGGATAGCGGAGCCTATCTCTCCTGTGGGGCGCTGTGGCTGTGCTTGT

1861   CGCTGGATGAGCAGCGGCGTAAAACGCCCCCTCAGGAAAGCGACTATACCCACTACGCCT

1921   TCAGCGTGGCGGAAGAAGAGTTTGCCGGGGTGGTGGCTCTGCTGGCGCAGGCGGGGGCTG

1981   AGGTATGGAAAGATAACCGCAGTGAAGGGGCGTCTTACTATTTTCTCGACCCTGACGGCC

2041   ATAAGCTGGAGCTGCATGTGGGGAATCTGGCGCAGCGGCTGGCCGCCTGTCGCGAACGCC

2101   CCTACAAGGGGATGGTCTTTTTTGATTGACGGGTTAGTTCAGCTTACTGCCGGATTTCAA

2161   CGTGCAGATCCACAGC
```

**Fig. S4.** (**A**) Genetic organisation of *exc_L, exc_C* and *fosA3* for synthesis. Putative promoter regions (consensus -35 and -10) are shown. The region shown by the blue bar has been found to be deleted in the commercially synthesised gBlock, supplied by IDT. (**B**) Nucleotide sequences of *exc(L)_exc(C)_fosA3* gBlock. Grey, green and blue shaded nucleotides are the coding region of *exc_L, exc_C* and *fosA3* genes, respectively.

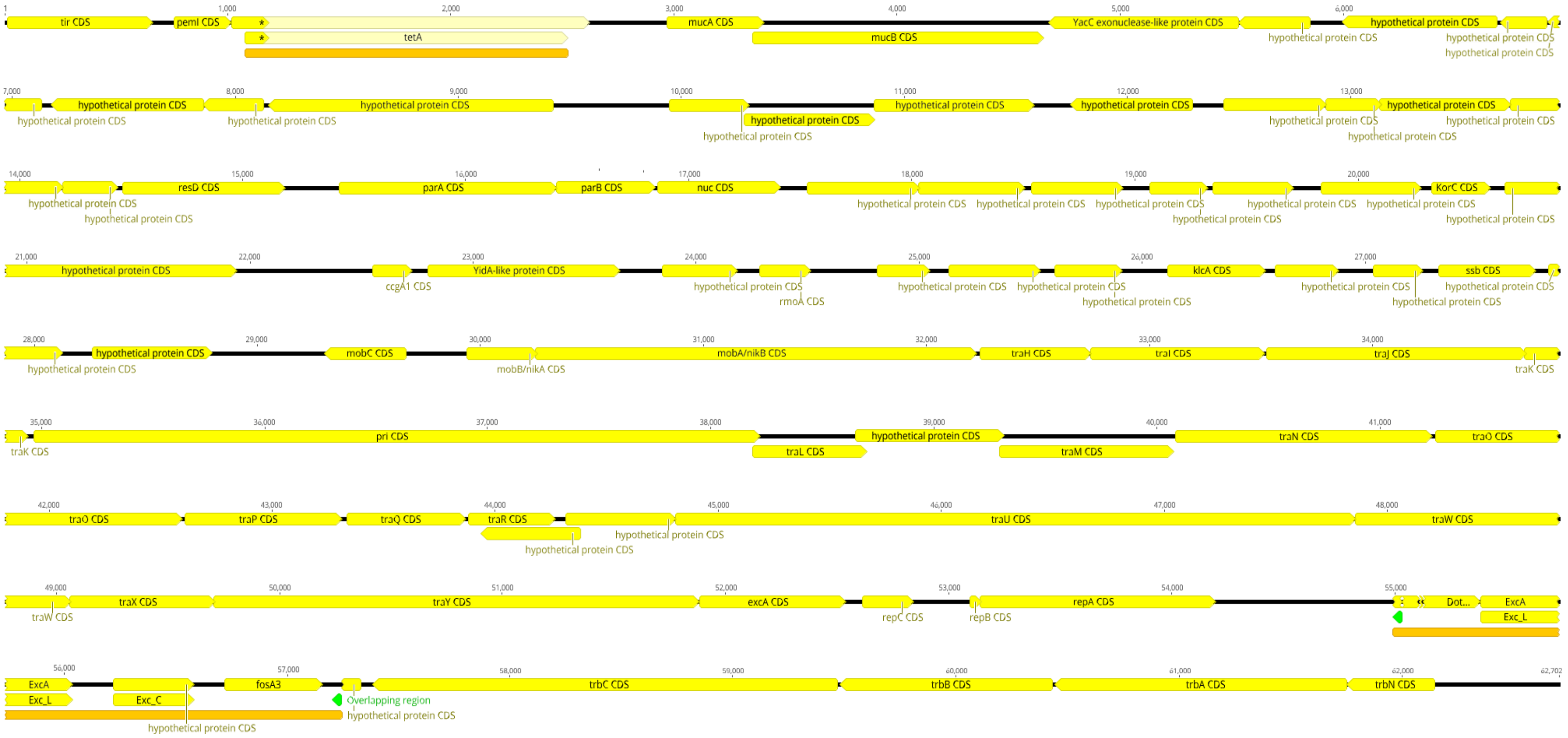


**Fig. S5.** The physical map of the conjugative probiotic plasmid PB1.1 (Size: 62702 bp). Two orange regions are for two independent insertion locations.

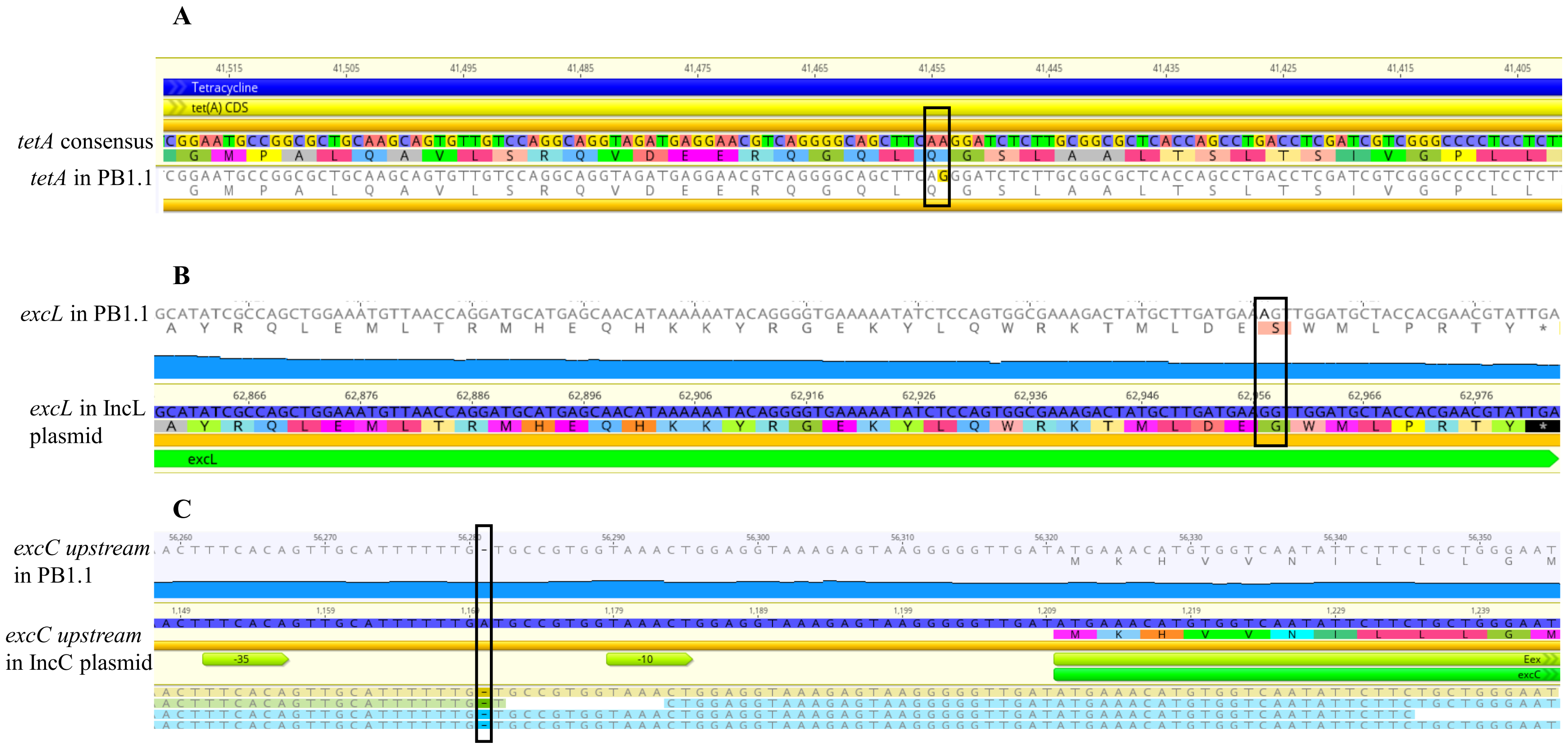


**Fig. S6.** Mutations identified in the constructed pPB1.1. Changed nucleotide and amino acid are marked with black boxes.
